# Interaction Between Ploidy, Breeding System, and Lineage Diversification

**DOI:** 10.1101/709329

**Authors:** Rosana Zenil-Ferguson, J. Gordon Burleigh, William A. Freyman, Boris Igić, Itay Mayrose, Emma E. Goldberg

## Abstract

If particular traits consistently affect rates of speciation and extinction, broad macroevolutionary patterns can be understood as consequences of selection at high levels of the biological hierarchy. Identifying traits associated with diversification rate differences is complicated by the wide variety of characters under consideration and the statistical challenges of testing for associations from comparative phylogenetic data. Ploidy (diploid vs. polyploid states) and breeding system (self-incompatible vs. self-compatible states) have been repeatedly suggested as possible drivers of differential diversification. We investigate the connections of these traits, including their interaction, to speciation and extinction rates in Solanaceae. We show that the effect of ploidy on diversification can be largely explained by its correlation with breeding system and that additional unknown factors, alongside breeding system, influence diversification rates. These results are largely robust to allowing for diploidization. Finally, we find that the most common evolutionary pathway to polyploidy in Solanaceae occurs via direct breakdown of self-incompatibility by whole genome duplication, rather than indirectly via breakdown followed by polyploidization.

---

> *“Among life history traits, reproductive characters that determine mating patterns are perhaps the most influential in governing macroevolution.”*
>
> — Barrett et al. (1996)

## Introduction

Species accumulate across the tree of life at different rates. One possible explanation for this phenomenon is that various traits differentially affect rates of diversification. Dramatic increases in phylogenetic and phenotypic data, along with methodological advances, have greatly accelerated the search for traits that influence diversification. Nevertheless, identifying focal traits associated with rates of speciation and extinction remains a challenge (e.g., Maddison and FitzJohn 2015; Rabosky and Goldberg 2015; Beaulieu and O’Meara 2016; Rabosky and Goldberg 2017). One difficulty is that speciation and extinction likely do not depend on a single character, so the biological and environmental contexts in which traits occur can lead to complex interactions that affect lineage diversification (Beaulieu and O’Meara 2016; Caetano *et al*. 2018; Herrera-Alsina *et al*. 2018). Consequently, examining the association of only one character with diversification patterns can be misleading. Here, we embrace this challenge by jointly investigating two characters thought to influence speciation and extinction rates—ploidy level and breeding system—while allowing for their interactions, and other confounding factors, to change diversification. We also test whether adding one more trait and increasing model complexity is worthwhile.

Polyploidization is a remarkably common mutation in plants (Husband *et al*. 2013; Zenil-Ferguson *et al*. 2017). The widespread variation in ploidy has long been considered a salient feature of flowering plant lineages (Stebbins 1938). An increase in ploidy can alter many traits and affect a variety of evolutionary and ecological processes (Ramsey and Schemske 2002; Sessa 2019). At shallow evolutionary time scale, polyploids were found to have an overall lower net diversification rate than diploids across many vascular plant clades (Mayrose *et al*. 2011, 2015). However, recent genomic studies have inferred numerous paleo-polyploidizations, including some preceding the emergence of highly diverse plant clades (Soltis *et al*. 2014; Landis *et al*. 2018), suggesting that whole genome duplications may have played an important role driving innovation and diversification in plants. Evidence of paleo-polyploidy within the genomes of diploid extant plants also implies pervasive diploidization, the return of polyploids to the diploid state, throughout the angiosperm phylogeny (Soltis *et al*. 2015; Dodsworth *et al*. 2015). Our analyses re-examine the association between ploidy and lineage diversification by extending the approach of Mayrose *et al*. (2011, 2015) to include transitions from polyploid to diploid states and potential unobserved factors affecting diversification patterns.

Breeding system shifts—changes in the collection of physiological and morphological traits that determine the likelihood that any two gametes unite—are also remarkably common and crucially affect the distribution and amount of genetic variation in populations (Barrett 2013). In particular, a variety of genetic self-incompatibility (SI) systems cause plants to reject their own pollen, and the loss of such mechanisms, yielding self-compatibility (SC), is a commonly observed transition in flowering plant evolution (Stebbins 1974; Igić *et al*. 2008). Previous analyses reported higher rates of diversification for SI than for SC lineages in Solanaceae (Goldberg *et al*. 2010). Similarly, heterostylous SI lineages in Primulaceae seem to diversify faster than SC lineages (de Vos *et al*. 2014), as do outcrossing lineages in Onagraceae (Freyman and Höhna 2019). Although these findings suggest a consistent macroevolutionary role of breeding system, it is unlikely to be the sole character determining lineage diversification. We investigate the relationship of breeding system to speciation and extinction rates in the context of ploidy and other unobserved factors.

Polyploidy and self-fertilization are widely thought to be associated (Stebbins 1950). Whole genome duplication may facilitate the transition to selfing by masking inbreeding depression, or self-fertilization may facilitate polyploidy establishment by by avoiding the lower fitness that characterize early-forming polyploids (Levin 1975; Ramsey and Schemske 1998; Barringer 2007; Barrett 2008; Husband *et al*. 2008).Additionally, in RNase-based gametophytic SI systems, polyploidization directly causes the loss of SI (Stout and Chandler 1942; Lewis 1947). In these systems, SI occurs because haploid self-pollen grains, with one S-allele at the locus controlling the SI response, are unable to detoxify the S-RNase produced by the same S-allele in the style (Kubo *et al*. 2010). The unreduced pollen of diploids, however, can contain two S-alleles expressed in pollen, which jointly gain the ability to detoxify the S-RNases produced by any maternal genotype (Entani *et al*. 1999; Tsukamoto *et al*. 2005; Kubo *et al*. 2010). Initial mutant individuals with pollen containing doubled haploid genomes are consequently capable of self-fertilization, with exceedingly few exceptions (Hauck *et al*. 2002; Nunes *et al*. 2006). RNase-based SI is regarded as ancestral in eudicots (Igić and Kohn 2001; Steinbachs and Holsinger 2002), and it is expressed in all SI species of Solanaceae examined to date. The absence of SI polyploids in this family yields a strong correlation between ploidy and breeding system (Robertson *et al*. 2011).

We address two macroevolutionary questions about the correlated evolution of ploidy and breeding system. First, we investigate their joint influence on rates of speciation and extinction. Each character alone has been shown to be associated with differential lineage diversification, but if their effects are not additive, studying them separately may not reveal their combined effect. Second, we examine the order of transitions in the two characters. Evolution commonly proceeds from diploid to polyploid, and from SI to SC states, but there are two paths by which diploid SI lineages can eventually become SC polyploids. Loss of SI in diploids could be directly caused by polyploidization (as explained above, for RNase-based SI systems), resulting in a one-step pathway to SC polyploids. Alternatively, SI diploids could first transition to SC without an increase in ploidy, and subsequently undergo polyploidization, resulting in a two-step pathway to SC polyploids. Robertson *et al*. (2011) compared the contributions of these two paths, finding that evolution from SI diploids to SC polyploids is more likely to proceed via the one-step pathway over short timescales, but via the two-step pathway over long timescales. They considered only transitions among the states, however, and we investigate whether these results hold true when allowing for differences in lineage diversification.

In the present study, we employ an extended framework of state speciation and extinction models, which simultaneously model transitions between the discrete states of a trait and different rates of speciation and extinction associated with each of those states (‘SSE’ models; Maddison *et al*. 2007; FitzJohn 2012). We start by fitting binary state speciation and extinction models to ploidy and breeding system independently (Maddison *et al*. 2007). We follow by fitting models that incorporate hidden states, which reduce the chance that the effect of the focal trait (ploidy or breeding system) on diversification is found to be significant when in reality, it is simply background heterogeneity in the diversification process what produces the diversification patterns in the phylogeny (Beaulieu and O’Meara 2016). We compare the proposed models against their character independent counterparts (Beaulieu and O’Meara 2016) to investigate whether ploidy or breeding system are actually affect the diversification process. Next, we model ploidy and breeding system jointly to assess their combined influence on diversification, with or without an additional hidden state. Using the ploidy and breeding system model without hidden states, we quantify the relative contributions of the two pathways from SI diploids to SC polyploids. We also aggregate states within these joint models of ploidy and breeding system in order to test whether increasing complexity from one trait to two traits significantly improves our understanding of the diversification process. Furthermore, we extend all the models involving ploidy to investigate the potential effects of including diploidization. Our results highlight the importance of considering non-additive effects of traits on net diversification rates under the presence of unobserved factors, in order to detect strong biologically-driven processes dictating the diversification patterns.

## Methods

### Data

Chromosome number data were obtained for all Solanaceae taxa in the Chromosome Counts Database (CCDB; Rice *et al*. 2015), and the ca. 14,000 records were curated semi-automatically using the CCDBcurator R package (Rivero *et al*. 2019). CCDB contains records from original sources that have multiple complex symbol patterns denoting multivalence, or irregularites of chromosome counts. After a first round of automatic cleaning, we examined results by hand and corrected records as necessary. Our hand-curated records were also contrasted against the ploidy dataset from Robertson *et al*. (2011), original references therein, and against ploidy data in the C-value DNA dataset from Bennett and Leitch (2005). By comparing three different sources of information, we were able to code taxa as diploid, *D*, or polyploid, *P*. For the majority of species, ploidy was assigned according to information from the original publications included in the C-value DNA dataset (Bennett and Leitch 2005). For taxa without ploidy information but with information about chromosome number, we assigned ploidy based on the multiplicity of chromosomes within the genus/family, or based on SI/SC classification. For example, *Solanum betaceum* did not have information about ploidy level but it has 2n=24 chromosomes, so since *x* = 12 is the base chromosome number of the genus *Solanum* (Olmstead and Bohs 2007), we assigned *S. betaceum* as diploid. Additionally, because of the absence of SI polyploids (explained above and below), species known to be SI could be scored as diploid. Species with more than one ploidy level were assigned the most frequent ploidy level recorded or the smallest ploidy in case of frequency ties.

Breeding system states were scored as self-incompatible, *I*, or self-compatible, *C*, based on results curated from the literature (as compiled in Igić *et al*. 2006; Goldberg *et al*. 2010; Robertson *et al*. 2011; 2012). Most species could unambiguously be coded as either *I* or *C* (Raduski *et al*. 2012). Following previous work, we coded any species with a functional SI system as *I*, even if SC or dioecy was also reported. Dioecious species without functional SI were coded as *C*.

Resolution of taxonomic synonymy followed Solanaceae Source (PBI *Solanum* Project 2012). Hybrids and cultivars were excluded because ploidy and breeding system can be affected by artificial selection during domestication. Following the reasoning outlined in Robertson *et al*. (2011), we closely examined the few species for which the merged ploidy and breeding system data indicated the presence of self-incompatible polyploids. Although SI populations frequently contain some SC individuals, and diploid populations frequently contain some polyploid individuals, in no case did we find convincing data for a naturally occurring SI polyploid population (discussed in Robertson *et al*. 2011). Because of the resulting absence of polyploid SI populations, as well as the functional explanation for polyploidy disabling gametophytic SI systems with non-self recognition (see the Introduction), we consider only three observed character states: self-incompatible diploids (*ID*), self-compatible diploids (*CD*), and self-compatible polyploids (*CP*).

Matching our character state data to the largest time-calibrated phylogeny of Solanaceae (Särkinen *et al*. 2013) yielded 651 species with ploidy and/or breeding system information on the tree. Of these, 368 had information for both states. The number of species in each combination of states is summarized in Fig. 1A. We retained all 651 species in each of the analyses below because pruning away tips lacking breeding system in the ploidy-only analyses (and vice versa) would discard data that could inform the diversification models. A total of 372 taxa without any information about breeding system or ploidy were excluded.

**Figure 1:**
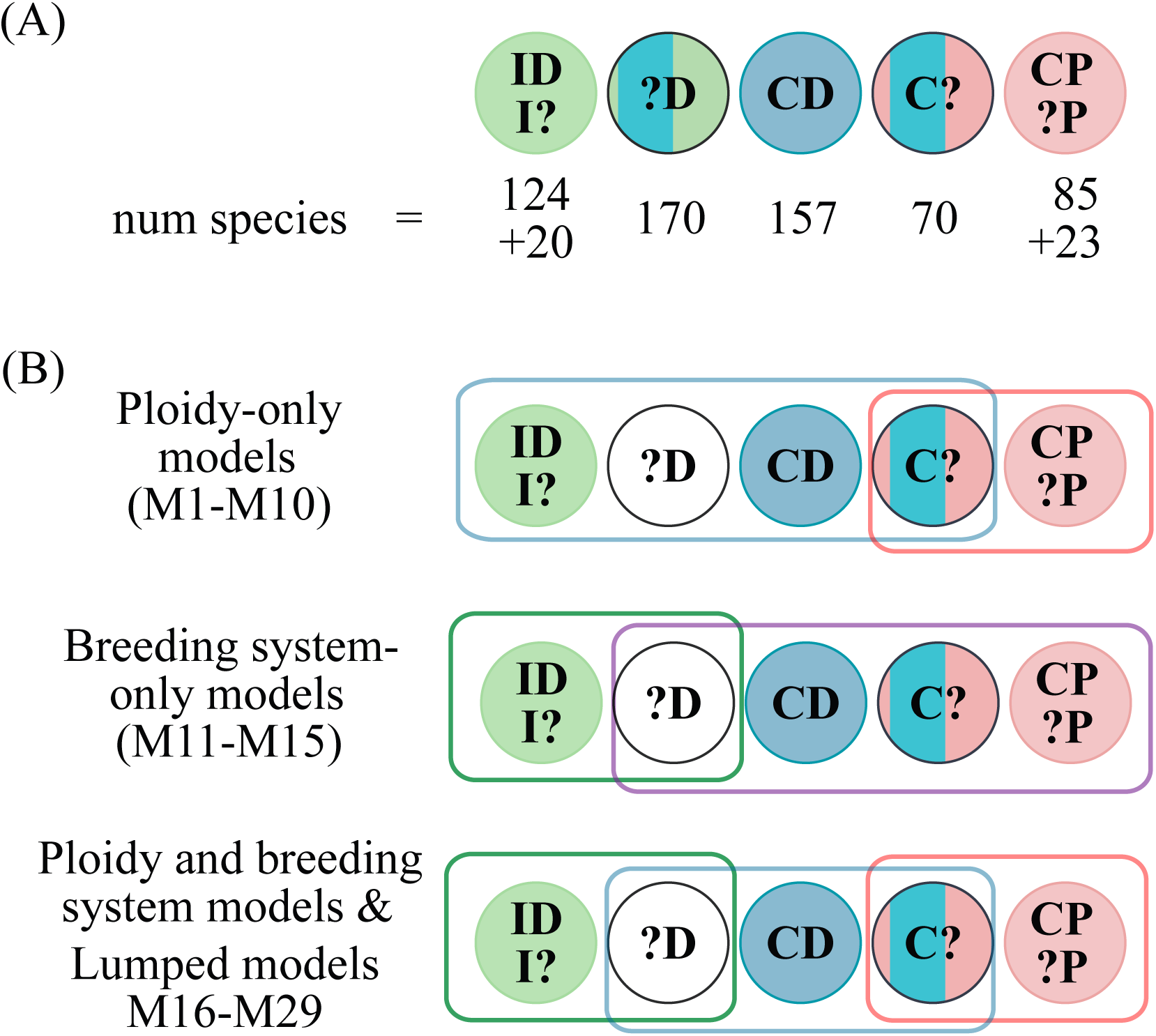
Character states used in the models. (A) Each species retained on the phylogeny belonged to one of five possible categories, depending on whether ploidy and/or breeding system were known. Number of species in each category is indicated; for example, 70 species are self-compatible with unknown ploidy. Character state abbreviations are: *I* for self-incompatible, *C* for self-compatible, *D* for diploid, *P* for polyploid, for unknown. Because polyploidization breaks this form of self-incompatibility, self-incompatible species with unobserved ploidy (*I*?) are taken to be diploid (*ID*), and polyploid species with unobserved breeding system (?*P*) are taken to be SC (*CP*). (B) Category groupings into states for each model class. In the ploidy-only models (M1-M10), states are coded as *D* & *P* when uncertain/consistent with either state; in the breeding system-only models (M11-M15) such states are coded as *C*; in the ploidy and breeding system models (M16-M29), they are coded as *CD* & *CP*. In the hidden-trait models, all species could take on either of two ‘hidden’ character states. Two species, *Lycium californicum* and *Solanum bulbocastanum*, are simultaneously *ID* and *CP*, and by adding them the sample adds to the total of 651 taxa used for analyses.

The Supplementary Information contains citations for the numerous original data sources. The Dryad archive contains the data and tree files used for analyses.

### Models

In order to test our hypotheses about lineage diversification and trait macroevolution, we fit 29 state-dependent speciation and extinction models (BiSSE, MuSSE, HiSSE; Maddison *et al*. 2007; FitzJohn 2012; Beaulieu and O’Meara 2016). SSE models contain parameters that describe per-lineage rates of speciation and extinction, specific to each character state (denoted *λ* and *µ*, respectively, with subscripts to indicate the state), along with rates of transitions between states (denoted *ρ* for polyploidization, *δ* for diploidization, and *q_IC_* for loss of self-incompatibility). The full set of models and all their rate parameters are detailed in Fig. S1. Here, we summarize how each model allows us to assess whether diversification is best explained by variation in ploidy, breeding system, their combination, or some unknown factor.

#### Ploidy and diversification

In order to investigate the association between ploidy level and diversification, we first employed a model (labeled M1), previously used by Mayrose *et al*. (2011), with each species classified as diploid (*D*) or polyploid (*P*). Although this model can be powerful in studies of trait evolution, it is prone to incorrectly reporting that a trait is associated with diversification differences (Maddison and FitzJohn 2015; Rabosky and Goldberg 2015). We therefore define several models that incorporate additional forms of diversification rate heterogeneity.

The second ploidy model (M2) includes a binary hidden trait that subdivides each observed state. In this trait-independent model known as CID (Beaulieu and O’Meara 2016), hidden traits can affect diversification but the observed traits do not. Comparing M1 and M2 allows us to test whether diversification rate heterogeneity is better explained by ploidy or by some unknown factor.

We additionally fit three models in which both ploidy and a hidden trait could influence diversification (M3–M5). These models differ in whether transitions between the hidden states are symmetric (M3) or asymmetric(M4), and whether the polyploidization rate depends on the hidden state(M5). Comparing M1– M5 allows us to test whether ploidy is associated with diversification differences on top of the differences potentially explained by an unknown factor.

We further fit the analogues of these five models but including a rate parameter *δ* for transitions from polyploid to diploid (M6–M10). These comparisons allow us to assess whether our conclusions about ploidy and diversification are robust to the possibility of diploidization.

#### Breeding system and diversification

We propose five breeding system models following the same logic as the ploidy models above. Under the simplest breeding system and diversification model (M11), species are classified as self-incompatible (*I*) or self-compatible (*C*). This is the same model as in the analysis presented in Goldberg *et al*. (2010) but with an updated phylogeny (Särkinen *et al*. 2013) and a larger aggregated dataset.

We then add models to allow diversification to be influenced by only a hidden trait (M12), or by both breeding system and a hidden trait (M13–M15, with varying degrees of complexity in the hidden trait transitions Table 2). Similar models were used by Freyman and Höhna (2019) to study diversification in Onagraceae.

**Table 1:** Bayes factors of ploidy-only models without diplodization. Numbers greater that 1 mean moderate support relative to the model in the row label. Conventional threshold for ‘strong’ support is 10. The best models are written in bold.

| Model | Marginal log-likelihood | M2 | M3 | M4 | M5 | Evidence |
| --- | --- | --- | --- | --- | --- | --- |
| M1. D/P | -1283.76 | 59.90 | 49.23 | 60.48 | 58.82 | Every model strongly preferred over M1 |
| **M2. CID D/P** | **-1223.86** |  | -10.66 | 0.579 | -1.079 | Strong preference for M2 over M1 |
| M3. D/P+A/B | -1234.52 |  |  | 11.24 | 9.58 | Asymmetric rates strongly preferred over symmetric |
| **M4. D/P+A/B asym** | **-1223.28** |  |  |  | -1.658 | Moderate evidence for only asymmetric hidden rates |
| M5. D/P+A/B all asym | -1224.93 |  |  |  |  |  |

**Table 2:** Bayes factors of breeding system-only models. Results indicate that the HiSSE models with asymmetric rates (M14, M15) are strongly preferred over every other model.

| Model | Marginal<br>log-likelihood | M12 | M13 | M14 | M15 | Evidence |
| --- | --- | --- | --- | --- | --- | --- |
| M11. I/C | -1309.07 | 41.13 | 38.59 | 61.34 | 61.40 | Every model strongly preferred over M11 |
| M12. CID I/C | -1267.93 |  | -2.53 | 20.21 | 20.37 | Models with asymmetric rates are preferred over M12 |
| M13. I/C+A/B | -1270.47 |  |  | 22.75 | 22.80 | Asymmetric rates strongly preferred over symmetric |
| **M14. I/C+A/B asym** | **-1247.72** |  |  |  | 0.05 | No evidence |
| **M15. I/C+A/B all asym** | **-1247.66** |  |  |  |  |  |

Self-incompatibility is homologous in all Solanaceae species in which S-alleles have been cloned and controlled crosses performed. All species sampled to date possess a non-self recognition, RNase-based gametophytic self-incompatibility (shared even with other euasterid families; Ramanauskas and Igić 2017). Furthermore, species that are distantly related within this family carry closely-related alleles, with deep trans-specific polymorphism at the locus that controls the SI response (Ioerger *et al*. 1990; Igić *et al*. 2006). Thus, there is strong evidence in Solanaceae that the *I* state is ancestral in the family, and that the SI mechanism was not regained. For all breeding system models, we estimated a transition rate from *I* to *C* but not the reverse (*q_CI_* = 0).

#### Ploidy, breeding system, and diversification

Ploidy and breeding system might influence lineage diversification individually, but these two traits also have an intricate association (discussed in the Introduction). Therefore, we considered several multi-state models that investigate the contribution of both traits and the allowable transitions between them.

The simplest model (M16) classifies each species as either SI diploid (*ID*), SC diploid (*CD*), or SC polyploid (*CP*); recall that SI polyploids do not occur. Each of these states may again be associated with different rates of speciation and extinction, and the allowable transitions are loss of SI within the diploid state (from *ID* to *CD*), loss of SI via polyploidization (from *ID* to *CP*), and polyploidization while SC (from *CD* to *CP*).

As for the previous models of only one trait, we then allow diversification to be influenced by only a hidden trait (M17), or by ploidy, breeding system, and a hidden trait (M18–M20) with varying degrees of complexity in the hidden trait transitions (similar to Caetano *et al*. 2018; Huang *et al*. 2018). We also fit the analogous models but allowing for diploidization (M21–M25).

#### Lumped models

The models described so far allow us to assess the contributions of our two focal characters—ploidy and breeding system—to lineage diversification, but they do not reveal whether it is valuable to include both characters in the analysis. To answer this question with statistical model comparisons requires comparing the likelihood of the data given each model. This is impossible for the ploidy and breeding system models presented so far, however, because the data are different for the different models: they use either the *D/P* or the *I/C* or the *ID/CD/CP* state spaces. Therefore, the use of different data results in incomparable models.

In order to compare fits of ploidy-only vs. breeding system-only vs. combined trait models, we use the technique of ‘lumping’ states together (Tarasov 2019). We use the state space of the *ID/CD/CP* model but constrain the rate parameters to mimic the behavior of the single-trait models. To lump states requires that the transition rates from the lumped state to the singular state be equal (Tarasov 2019). First we lump together *ID* and *CD* to form the diploid state, mimicking the *D/P* model (M26). Proposing a lumped ploidy model by aggregating *ID* and *CD* requires forcing the rate of polyploidization from *ID* and *CD* to *CP* to be equal (i.e. *ρ*_0_ = *ρ_I_*= *ρ_c_*), but also requires assuming that the rates of speciation and extinciton for *ID* and *CD* to be equal. Therefore, we define the new parameters *λ_D_*and *µ_D_* that are the same for each of the two diploid states *ID* and *CD*.We used the same procedure to lump together *CD* and *CP* to form the self-compatible state, mimicking the *I/C* model (M28). In this particular case, the rate from *CD* to *CP* back to *ID* is zero and equal for both, so the model is lumpable. However, to fully mimic the breeding system model, we assume that the rate of selfing is equal (i.e. *q*_0_ = *q_IC_* = *rho_I_*) and the rates of speciation and extinction for both *CD* and *CP* are the same (new parameters *λ_C_* and *µ_C_*). We further add a hidden character to each of these models (M27 and M29), and then compare this group of models (Table 3).

**Table 3:** Test for adding a focal trait to a binary state model via Bayes factors. Three-state models are moderately preferred over two-state models. The evidence is stronger towards the inclusion of breeding systems in ploidy models (M26, M27) than the inclusion of ploidy in breeding system models (M29, M29)

| Model | Marginal<br>log-likelihood | Comparison | $K=\ln(\text{BF}(M_i, M_{ii}))$ | Preferred<br>Model (Evidence) |
| --- | --- | --- | --- | --- |
| M26. Lumped D/P<br>**M16. ID/CD/CP** | -1463.22<br>**-1459.11** | M26 vs. M16 | 4.10 | M16 (Moderate) |
| M27. Lumped D/P+A/B<br>**M23. ID/CD/CP +A/B** | -1417.67<br>**-1414.00** | M27 vs. M23 | 3.678 | M23 (Moderate) |
| M28. Lumped I/C<br>**M16. ID/CD/CP** | -1458.41<br>**-1459.11** | M28 vs. M16 | -0.69 | No evidence |
| M29. Lumped I/C+ A/B<br>**M23. IC/CD/CP+A/B** | -1416.60<br>**-1414.00** | M29 vs. M23 | 2.60 | M23 (Moderate) |

We do not include additional models with diploidization because this reverse ploidy transition renders the models non-lumpable. When including diploidization, transitions from *CP* to *CD* are at rate *δ* but transitions from *CP* to *ID* do not occur. Because *ID* and *CD* would be lumped to mimic the *D/P* model, this model is non-lumpable when *δ /*= 0. Thus, we can compare models to test whether it is advantageous to include both traits, but only when ignoring diploidization.

#### Pathways to polyploidy

Considering ploidy and breeding system together, there are two evolutionary pathways from SI diploid to SC polyploid (Brunet and Liston 2001; Robertson *et al*. 2011). In the one-step pathway, the *CP* state is produced directly from the *ID* state when whole genome duplication disables SI. In the two-step pathway, the *CD* state is an intermediate: SI is first lost, and later the SC diploid undergoes polyploidization. We quantify the relative contribution of these pathways to polyploidy in two ways, each using the median estimates of rates from the simplest model that includes both traits (M16). Our results will differ from those of Robertson *et al*. (2011) because our transition rate estimates come from a dated phylogeny and a model that allows for state-dependent diversification.

Both of our methods are based on a propagation matrix that describes flow from *ID* to *CP*, as in Robertson *et al*. (2011). We insert an artificial division in the *CP* state, so that one sub-state contains the *CP* species that arrived via the one-step pathway and the other substate contains the *CP* species that arrived via the two-step pathway. We consider unidirectional change along each step of the pathway in order to separate them into clear alternatives, and because in this family there is no support for regain of SI, and no strong support for diploidization (see below).

First, we consider only the rates of transitions between these states, placing them in the propagation matrix Q. The matrix P = exp(Q*t*) then provides the probabilities of changing from one state to any other state after time *t*. Closed-form solutions for the two pathway probabilities are provided in Robertson *et al*. (2011).

Second, we consider not only transitions between states but also diversification within each state. State-dependent diversification can change the relative contributions of the two pathways. For example, if the net diversification rate is small for *CD*, the two-step pathway will contribute relatively less. We therefore include the difference between speciation and extinction along the diagonal elements of the propagation matrix. As before, matrix exponentiation provides the relative chance of changing from one state to any other state after time *t*. The calculations of the propagation matrix are not probabilities because diversification changes the number of lineages as time passes. We can still use ratios, however, to consider the relative contribution of each pathway, analogous to the normalized age structure in a growing population (Leslie 1945).

### Statistical inference

#### Model fitting

Parameters for each of the 29 models were coded as graphical models and Bayesian statistical inference was performed with RevBayes (Höhna *et al*. 2016). Scripts for analyses and key results are available as Supplementary Information. We accounted for incomplete sampling in all analyses by setting the probability of sampling a species at the present to 651*/*3000 (using the method of FitzJohn *et al*. 2009) since the Solanaceae family has approximately 3,000 species (PBI *Solanum* Project 2012). For all models, we assumed that speciation and extinction parameters had log-normal prior distributions with means equal to the expected net diversification rate (number of taxa*/*[2 *×* root age]) and standard deviation 0.5. Priors for parameters defining trait changes were assumed to be gamma distributed with parameters *k* = 0.5 and *θ* = 1. For each model, a Markov chain Monte Carlo (MCMC; Metropolis *et al*. 1953; Hastings 1970) was run in the high-performance computational cluster at the Minnesota Supercomputing Institute, which allowed for 5,000 generations of burn-in and a minimum of 200,000 generations of MCMC for each of the models. Convergence and mixing of each MCMC chain was determined by ensuring the effective sample size of each parameter was over 200.

We report posterior distributions for the model parameters in Fig. 2, and also for the compound parameters of net diversification (*λ − µ*) and extinction fraction (*µ/λ*) for each state. Additionally, ancestral states at each node in the phylogeny were sampled jointly during the MCMC analyses every 100 generations. Ancestral state estimations for all models show the maximum *a posteriori* estimates of the marginal probability distributions for each of the 650 internal nodes for each of the models in Fig. 2. (Figs. S6, S8, S10, S12, S14 and S16).

**Figure 2:**
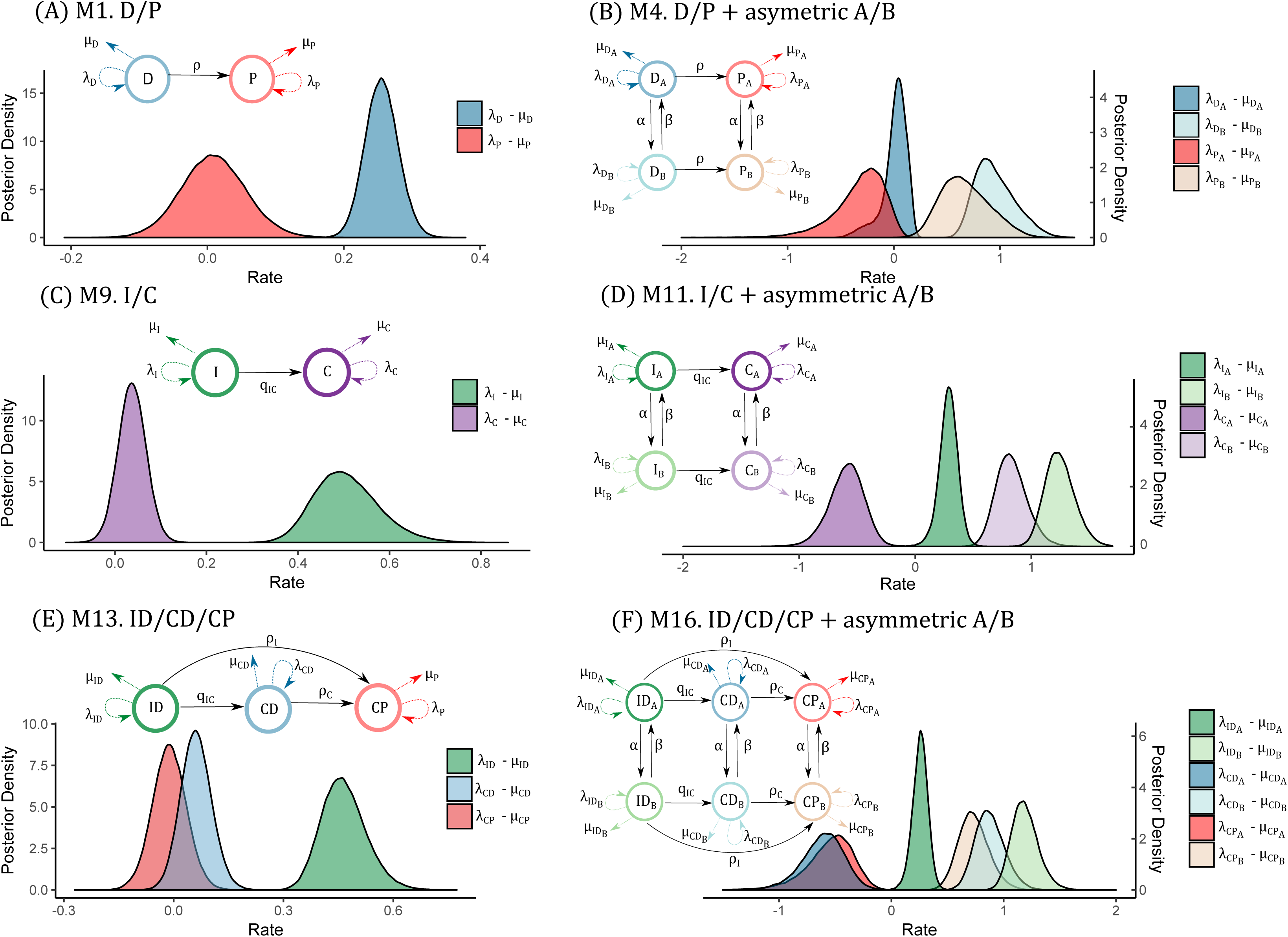
Net diversification rates for SSE models of focal traits with or without hidden state. Each panel contains a graphical summary of estimated model parameters and displays posterior distributions for net diversification estimates.

#### Model selection key questions

We calculated the marginal log-likelihood for each of the models using fifty stepping stone steps under the methodology of Xie *et al*. (2010), implemented in RevBayes (Höhna *et al*. 2016). Each stepping stone step was found by calculating at least 500 generations of burn-in followed by a total of 1,000 MCMC steps (Table 1).

Using the marginal likelihood values, we calculated Bayes factors to answer five key biological and methodological questions:

1. Are diversification patterns only determined by hidden states and not the traits of interest?— Comparison of character independent models against hidden state (Tables 1 and 2 and Tables S1–S3).
2. Are hidden states necessary to explain diversification rate heterogeneity?—Comparison of simple models against hidden state models (Table S4).
3. Does a second focal trait add information about the diversification process?—Comparison of lumped models against IC/CD/CP models (Fig. 3 and Table 3).
4. Are conclusions robust to assumptions about hidden state transitions?—Comparison amongst hidden states models with equal hidden rates and asymmetrical rates (Fig. S5 and Table S6).
5. Is there evidence for diploidization?—Comparison amongst models with and without diploidization (Fig. S2 and Table S5).

**Figure 3:**
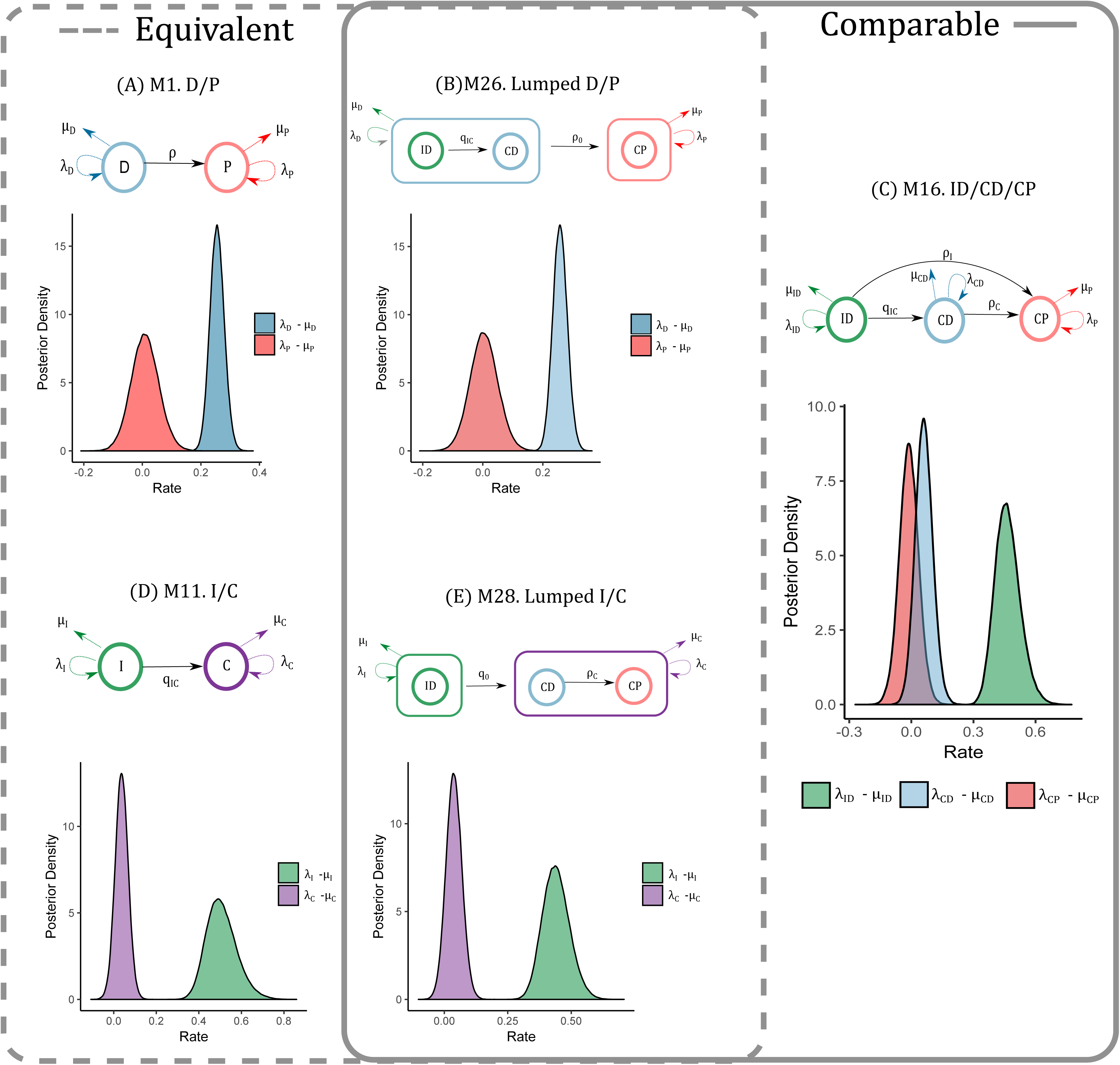
Comparing two and three state models is possible via lumpable models (M26, M28). These models used the same classifications as the three-state model M16 which allow for comparisons via Bayes Factors as shown in Table 3.

Each model comparison is reported with a Bayes factor on the natural log scale: the comparison between models *M*_0_ and *M*_1_ is *K* = *ln*(*BF*(*M*_1_*, M*_0_)) = ln[*P*(**X***|M*_1_) −*P*(**X***|M*_0_)]. There is ‘strong evidence’ for *M*_1_ when this value is more than 10, moderate support if the value is more than 1, and no evidence if the value is between −1 and 1. If the value of *K* is negative the evidence goes towards *M*_0_ (Kass and Raftery 1995).

## Results

### Trait-dependent diversification

#### Ploidy only

When considering ploidy alone, we found a larger net diversification rate for diploids than for polyploids, in agreement with Mayrose *et al*. (2011, 2015). This result holds with (model M1, Fig. 2A) or without the diploidization parameter (M6, Fig. S2A). Incorporating a hidden state in this model, however, reduces the clear separation in diversification rate estimates between diploids and polyploids (M4, Fig. 2B; M9, Fig. S2B). Statistical model comparisons show a clear preference for models in which only a hidden state affects diversification or a hidden state as well as ploidy (M2 and M4; Table 1). Results are similar when diploidization is included (Table S1). Thus, when other traits are ignored, the role of ploidy in net diversification is unclear, with marginal support for diploids having higher diversification rates, but rate differences perhaps better explained by another factor.

#### Breeding system only

When considering breeding system alone (M11, Fig. 2C), we found a larger net diversification rate for SI than for SC species, in agreement with Goldberg *et al*. (2010). When a hidden state is included, the large net diversification rate difference persists for one hidden state but is diminished for the other (M14, Fig. 2D). In the statistical model comparisons, the best two supported models include diversification differences due to both breeding system and to a hidden trait (M14 and M15, Table 2). Breeding system seems to play a role in diversification differences, though a hidden factor does as well.

#### Ploidy and breeding system together

When considering ploidy and breeding system together, the net diversification rate for SI diploids was greater than for either SC diploids or SC polyploids, with or without diploidization (M16, Fig. 2E; M21, Fig. S2E). Thus, the difference in net diversification associated with breeding system persists even when ploidy is included in the model. The reverse is not true: the association of ploidy with net diversification in the simplest ploidy-only model (M1, Fig. 2A, Fig. S2A) appears to be driven by the subset of diploids that are SI, while among SC species, net diversification rates for diploids and polyploids are similar.

When a hidden state is included, the separation in net diversification rate of ID vs. CD and CP persists within one hidden state but is reduced in the other (M19, Fig. 2F). The same general pattern remains when diploidization is included (Fig. S2F). Model comparisons clearly favor models that include ploidy, breeding system, and the hidden trait, against the character-independent model in which the focal traits do not influence diversification (Table S2; Table S3 with diploidization).

Using the lumped models, we find moderate support for obtaining a significantly better fit by adding breeding system to the ploidy-only model (M26 vs. M16, Table 3, Fig. 3BC). This is also true when a hidden trait is included (M27 vs. M23, Table 3, Fig. S3EF). A similar comparison in which ploidy is added to the breeding system-only model shows no preference for the model that also includes ploidy (M28 vs. M16, Table 3, Fig. 3CE). When including a hidden state, however, the model with both focal traits is moderately preferred over the model with only breeding system (M29 vs. M18, Table 3, Fig. S4EF).

From all of these types of statistical evidence, we conclude that breeding system (and a hidden factor) are strongly associated with diversification differences, and that ploidy plays a smaller role.

### Key questions about diversification and transitions

The above results include several of our statistical model comparison findings. Here we return to the five specific questions we targeted with our model comparisons.

1. Are diversification patterns only determined by hidden states and not the traits of interest? No, our focal traits are supported as having associations with diversification differences. In most cases, we find moderate to strong preference for models with the focal traits as well as hidden states, over models with only hidden states (Tables 1 and 2 and Tables S1–S3).
2. Are hidden states necessary to explain diversification rate heterogeneity? Yes, models with hidden states that influence diversification are strongly preferred over models containing only the focal traits (Table S4). This means that there are potentially many factors underlying diversification shifts within the family.
3. Does a second focal trait add information about the diversification process? Yes, in most cases models with both ploidy and breeding system are preferred over models with only one of the focal traits (Table 3 and Figs. S3 and S4).
4. Are conclusions robust to assumptions about hidden state transitions? Yes, we found that allowing different types of asymmetry in transitions within and between hidden states did not change our conclusions about net diversification differences. Hidden state models with asymmetric rates are, however, strongly preferred over models with equal rates between hidden states (Table S6), and they show stronger differences between some net diversification rates (Fig. S5). The effect of the asymmetry of the hidden rate transitions is better observed in the ancestral state estimations (Figs. S8, S12 and S16), which show that moving out of state A (dark colors) happens quickly with rate *α*, whereas evolving out of hidden state B (light colors) is slow with rate *β*.
5. Is there evidence for diploidization? Perhaps: when comparing models with diploidization against models without it, we found moderate evidence that models containing diploidization are preferred (Table S5). We discuss later some further challenges in identifying diploidization. We further found that our main conclusions about net diversification differences are not dependent on whether diploidization is included (Fig. S2).

### Pathways to polyploidy

There are two pathways by which SI diploid lineages eventually—given enough time—become SC polyploids. In the one-step pathway, polyploidization directly disables SI. In the two-step pathway, SI is first lost within the diploid state, followed by polyploidization. Determining the relative contribution of these pathways based on our transition rate estimates (median transition rate values from M16), we find that the one-step pathway is more likely on short timescales and the two-step pathway is more likely on long timescales (Fig. 4, left panels). Beginning with a single SI diploid lineage, when not much time has elapsed, the one-step pathway is more likely because it only necessitates a single event to reach the SC polyploid state. When more time has elapsed, the two-step pathway is more likely because the rate of loss of SI within diploids, *q_IC_*, is greater than the rate of polyploidization for SI species, *ρ_I_* (Fig. S15). That is, an *ID* lineage is more likely to begin its path to polyploidy with a transition to *CD*, but completing this path to *CP* takes longer. Robertson *et al*. (2011) reached the same conclusion. Our result is qualitatively unchanged when using transition rate estimates from the model that does not allow diversification differences related to the observed states (M17).

**Figure 4:**
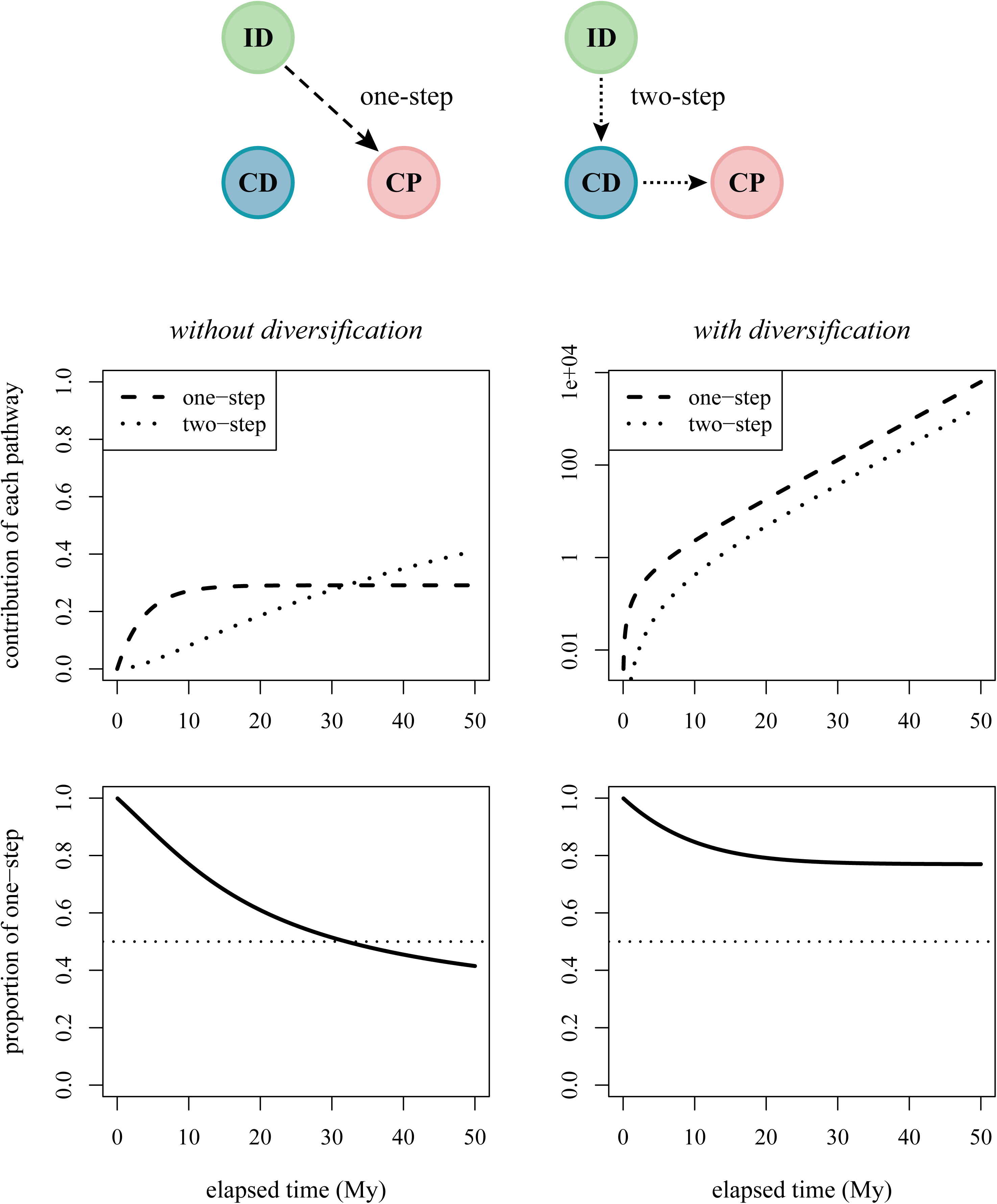
Contributions of the two pathways to polyploidy. The one-step pathway is direct ID*→*CP transitions. The two-step pathway consists of ID*→*CD*→*CP transitions. When considering only rates of transitions among the states (ignoring the diversification rate parameters), the one-step pathway dominates on short timescales and the two-step on long timescales (left panels). When also considering diversification within each state, the one-step pathway, in which polyploidization breaks down SI, dominates over any timescale (right panels). The top panels show the separate contributions of each pathway. The bottom panels show the proportional contribution of the one-step pathway (i.e., one-step / [one-step + two-step]).

The preceding conclusions, however, ignore the changes in numbers of lineages in each state due to speciation and extinction. By analogy, envisioning the states as stepping stones, the extent to which each stone grows or shrinks over time affects the utility of each possible path. Allowing different net diversification rates for each state (again using median rate estimates from M16), we find a qualitative difference in the relative pathway contributions. The lower rate of net diversification in the *CD* state, relative to *ID*, means that relatively fewer lineages are available to complete the second step of the two-step pathway, ending in *CP*. Consequently, even over long timescales, we find that the two-step pathway contributes less to the formation of polyploids (Fig. 4, right panels) when considering diversification as well as transitions.

## Discussion

Species are composed of vast assemblages of variable traits. Many traits are both heritable and possibly affect the propensity of species to perish or multiply (Lewontin 1970). Examining the effects of complex trait combinations on lineage diversification, however, remains challenging. Focusing first on ploidy and then on breeding system, we found that considering each trait in isolation provides an incomplete story. Considering them together, and in conjunction with another hidden factor, provides a more complete picture of macroevolutionary dynamics within Solanaceae. We hope our work serves as an example of how phylogenetic comparative methods can be used to disentangle the contributions of interacting traits to heterogeneous lineage diversification, and how to statistically argue for increasing complexity in diversification modeling.

### Interacting traits and lineage diversification

Previous analyses of the effects of ploidy on diversification found that diploids are associated with greater net diversification rates than polyploids across many angiosperm clades (Mayrose *et al*. 2011, 2015). We obtain a similar outcome when examining ploidy alone in Solanaceae (Fig. 2A), but a consistent effect of ploidy on diversification is not clear when we incorporate a hidden factor linked to diversification (Fig. 2B). Previous analyses of breeding system in this family found that SI may cause higher diversification rates, compared with SC (Goldberg *et al*. 2010; Fig. 2C). Our analyses that include a hidden trait recover the same pattern, with one important difference. On the background of one hidden state, we recover a net diversification rate for SC species that is positive and greater than the diversification rate of SI species on the background of the other hidden state (Fig. 2D). Therefore, SC may not be a ‘dead end’ when coupled with some unknown trait combinations or processes not modeled by these two traits. Our analyses also reveal that models of joint evolution of ploidy and breeding system are statistically preferred, and hint at how various trait combinations may be linked with diversification. We find that the highest net diversification rate is associated with SI diploids, while SC diploids have a lower diversification rate that overlaps with the net diversification of SC polyploids (Fig. 2E). Thus, breeding system appears to provide a relatively better explanation of diversification rate differences, with ploidy providing a secondary effect within SC species.

Throughout our numerous model comparisons, we find that inclusion of hidden states provides a considerably better fit (Table S4). This is consistent with the expectation that many processes, beyond those associated with the focal traits, can affect inference of speciation and extinction. It is, however, unclear exactly which processes are captured by the hidden states. For example, our results show that, to a varying extent, breeding system functions as a hidden state in the ploidy-centered analysis, and vice versa (Figs. S3 and S4). But the strong statistical support for processes not well explained by ploidy and breeding system (Table S4) tempts one to interpret the remaining variation as the effect of other measurable traits. For example, our data appear to show a rapidly-diversifying Australasian clade of mostly SC species within *Solanum*, which suggests that geography may play a role. Nevertheless, it is also possible that the addition of hidden states instead explains variation stemming from any of a number of unrelated processes or methodological artifacts, as previously discussed by Beaulieu and O’Meara (2016). In the absence of additional information, the hidden states can be viewed as a statistical trick, providing an easy way to model extra heterogeneity without directly representing a specific trait.

As more trait information becomes available for macroevolutionary studies, it is not only important to recognize the role of hidden states as a part of a general modeling approach, but also to question whether adding more traits to diversification studies is justified. More generally, we find that lumped models are useful for assessing the value of adding additional traits to already complex diversification models(Fig. 3 and Table 3).

Although we fit an extensive set of models in order to relax a variety of assumptions, we did not explore the process of trait change in conjunction with speciation. That is, our models all assume anagenetic trait evolution and ignore cladogenetic shifts. Anagenetic and cladogenetic changes can be separated with phylogenetic models (Mayrose *et al*. 2011; Goldberg and Igić 2012; Magnuson-Ford and Otto 2012). These have been applied to estimate the relative contribution of anagenetic and cladogenetic shifts in breeding system (Goldberg and Igić 2012) and polyploidization (Zhan *et al*. 2016; Freyman and Höhna 2019). We did not explore cladogenetic trait change because they would greatly increase in state space and model number of our already complex and extensive modeling framework. Although Goldberg and Igić (2012) found that allowing cladogenetic changes did not substantially affect inference of net diversification rates associated with breeding system, future work could test whether this process affects diversification rate estimates with the more complex state and parameter spaces of our other models.

### Pathways to polyploidy

With evolution predominantly in the direction from diploid to polyploid, and from SI to SC, surviving lineages will tend to become SC polyploids. We find that in Solanaceae, the pathway to this state is more likely to consist of a single step (*ID → CP*) than two steps (*ID → CD → CP*; Fig. 4). Although this question focuses on the process of state transitions, we also show that its answer is affected by the process of lineage diversification. We used a simple mathematical approach to investigate the contributions of the two pathways, but future work could instead rely on stochastic character mapping to estimate the numbers of each type of transition more directly.

Macroevolutionary transition rates represent a combination of time spent waiting for individuals with a new character state, and for that new state to become widespread within the species. For our traits, this consists of mutations that break SI or generate polyploid individuals, and selective pressures that cause fixation (or loss) of these mutants. Estimates of mutation rates are highly uncertain, but the chance of breakdown of SI within diploids is perhaps 10*^−^*^5^ per pollen grain; this includes breakdown by autopolyploidization (Lewis 1979), which is by itself estimated to occur approximately within the same order of magnitude (Ramsey and Schemske 1998). In contrast, we infer a macroevolutionary transition rate from *ID* to *CP* that is 2. 5 times greater than the rate from *ID* to *CD*, indicating that selection restricts the fixation of new polyploids more than of new SC mutants (Fig. 4, Robertson *et al*. 2011).

Our findings prompt several further questions about the macroevolutionary pathways of ploidy and breeding system. First, our support for the direct pathway is consistent with the idea that breakdown of SI by whole genome duplication—via diploid ‘heteroallelic’ pollen—may trigger the evolution of gender dimorphism as a different mechanism of inbreeding avoidance (Miller and Venable 2000). A further test of this hypothesis would additionally examine the propensity of polyploids generated through either pathway to become dioecious (Robertson *et al*. 2011). Second, we might wonder whether the propensity for a polyploid species to diversify depends on whether it arose via the one-step or two-step pathway. This could be tested with a different form of a hidden state model, in which the polyploid state is subdivided into parts, reflecting the arriving path taken to that state. Such an estimate of path-dependence could also include the possibility of different diversification rates for the subdivided *CP* substates. Third, the generality of our findings in other families remains to be assessed. An identical procedure could be used in other families with gametophytic SI. In clades with sporophytic SI systems, however, SI is not disabled by whole genome duplication, so there is no one-step pathway (Miller and Venable 2000; Mable 2004). The correlation between breeding system and ploidy may therefore be different in sporophytic systems, and it is unclear whether one of the two-step pathways might predominate.

### Diploidization

Polyploidization is known to be common in plants, but the pace and relative frequency of the reverse process—diploidization—remains unclear, and it is under active investigation (Dodsworth *et al*. 2015; Mandakova and Lysak 2018). Ignoring diploidization, if it is common, could cause underestimation and increase of uncertainty in the polyploidization rate. Therefore, we included a slate of models with a diploidization parameter, and show that our main conclusions are robust to this process (Fig. S2). These models also suggest modest statistical support for diploidization (Table S5), although our estimates of its rate were highly uncertain. Furthermore, additional lines of evidence for classifying species as diploid or polyploid (beyond the genus-level chromosome multiplicity that we primarily relied on) are needed for more reliable conclusions.

Other lines of evidence about the prevalence of diploidization within Solanaceae or its ancestors are mixed or even conflicting. On the one hand, polyploidy may have occurred prior to the origin of Solanaceae, rendering all extant ‘diploids’ secondarily derived. Ku *et al*. (2000) and Blanc and Wolfe (2004) posited that the lineage leading to cultivated tomato, *Solanum lycopersicum*, may have experienced one or more whole genome duplications. A subsequent analysis of synteny between grape and *Solanum* genomes, as well as genetic distances (Ks) between inferred paralogs within genomes of *Solanum* (tomato and potato), each suggested that this lineage experienced a likely round of ancient genome duplication or triplication (71 *±* 19 My; Tomato Genome Consortium 2012) likely pre-dating the origin of the family (49 *±* 3 My; Särkinen *et al*. 2013).

On the other hand, there is little evidence for the occurrence of diploidization after the origin of Solanaceae itself. Studies comparing genetic map-based genome synteny within a number of species in this family find no evidence for diploidization (Wu and Tanksley 2010). Instead, simple genome re-arrangements appear sufficient to explain chromosomal evolution within species in the cytogenetically conserved ‘x=12’ group, which includes tomato, potato, eggplant, pepper, and tobacco. Recovery of comparatively few rearrangements would require outstanding convergent loss of duplicated segments. Furthermore, whole genome duplication in a eudicot lineage ancestral to Solanaceae would clash with the evidence that the homologous mechanism of SI, which has been present continually in many families (Igić *et al*. 2006), breaks down nearly invariably in natural and induced tetraploids (Stone 2002; McClure 2009). Most problematically in this context, it is unclear how to explain the maintenance of trans-generic polymorphism at the orthologous S-loci in Solanaceae and other families if SI was previously broken down by polyploidization. Regardless of whether genome polyploidization, followed by widespread diploidization, is a dominant mechanism of genome evolution in Solanaceae, it is clear that more work is needed for a complete understanding of the joint evolution of ploidy and breeding systems.

## Conclusion

Heterogeneity in lineage diversification across time and clades is the rule, rather than the exception. This background heterogeneity makes it difficult to test for the association of any one, isolated trait with different rates of speciation or extinction. Our study provides an example of how diversification linked to a particular trait can be better assessed by a suite of more inclusive models that allow for alternative explanations— whether other traits or unknown factors. Additionally, our analysis of evolutionary pathways to polyploidy shows the importance of including diversification effects even when addressing questions that focus on trait evolution. Finally, although a growing number of plant traits have been studied to date, breeding systems indeed seem to be among the most influential in governing macroevolution (Barrett *et al*. 1996).

## Acknowledgements

Special thanks to Bob Thomson for donating computational resources that made possible testing the convergence of the models. We also thank Sergei Tarasov for clarifying lumpability concepts, and Carrie Tribble for troubleshooting some ancestral estimate figures. The computing resources were provided by the Minnesota Supercomputing Institute (MSI) at the University of Minnesota. This work was supported by the National Science Foundation grant DEB-1655478 (to EEG and IM), DEB-1655692 and NESCent sabbatical scholar award EF-0905606 (to BI), and the Israel Science Foundation 961/17 (to IM). We thank Dan Schoen and three anonymous reviewers for providing uncommonly thoughtful and helpful reviews, as well as Spencer Barrett for continually inspiring our work.

---

**Figure S1:**
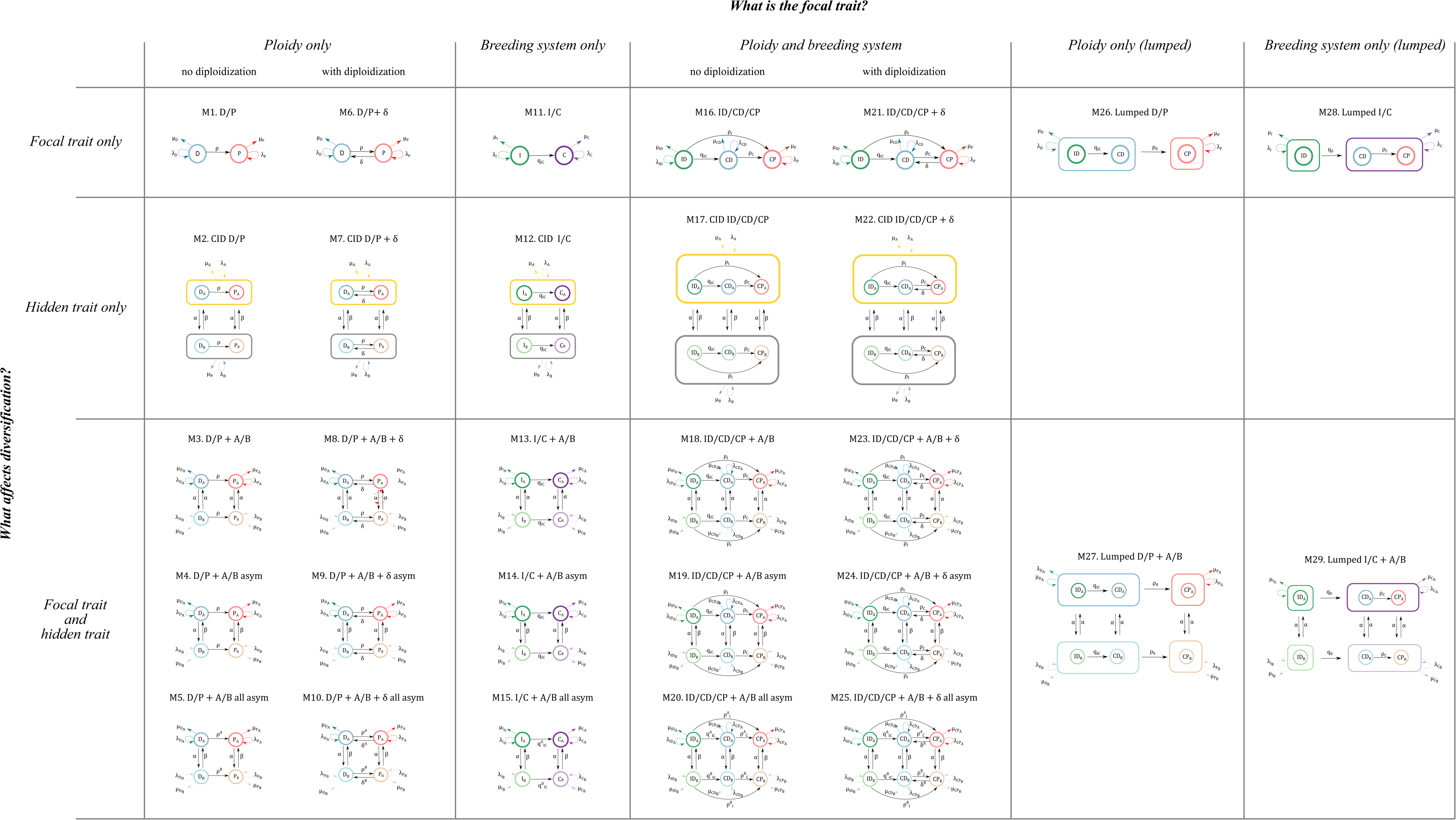
All models. [This figure is provided as a separate, large-format page.]

**Figure S2:**
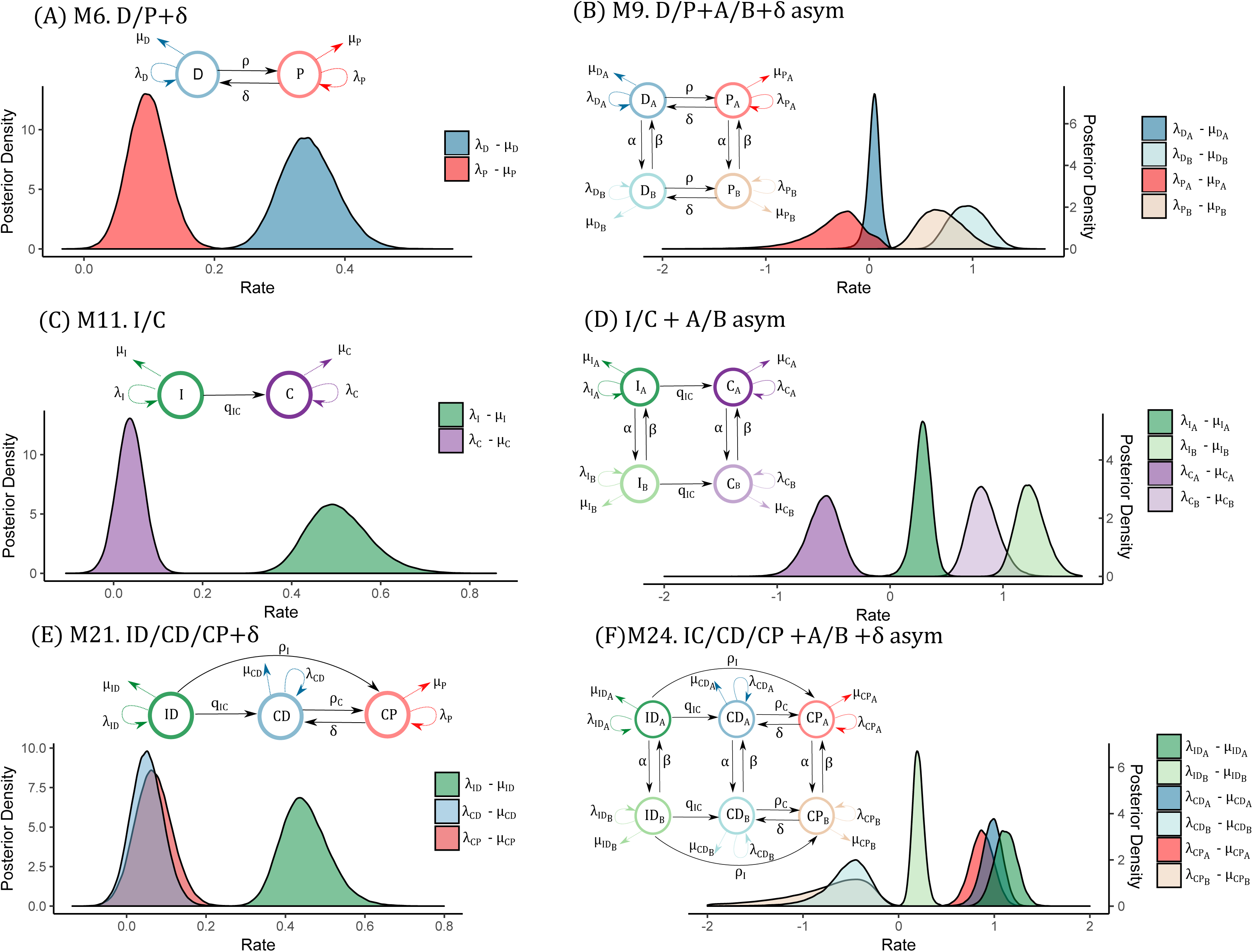
Posterior distribution for all the best models with diploidization

**Figure S3:**
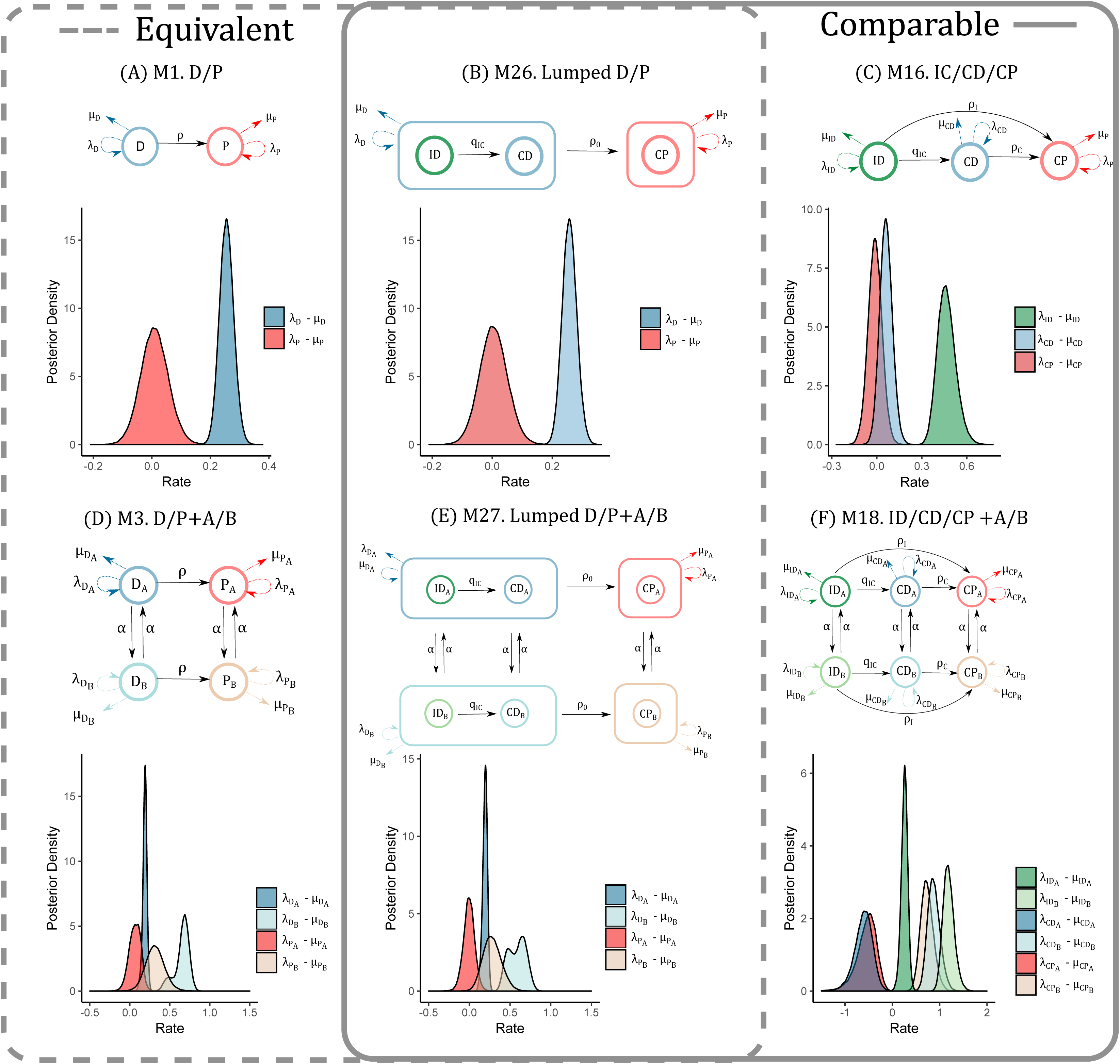
Lumped ploidy models to assess the effect of adding breeding system to create a three-state model. Moderate evidence exists that adding a breeding system is necessary Table 3

**Figure S4:**
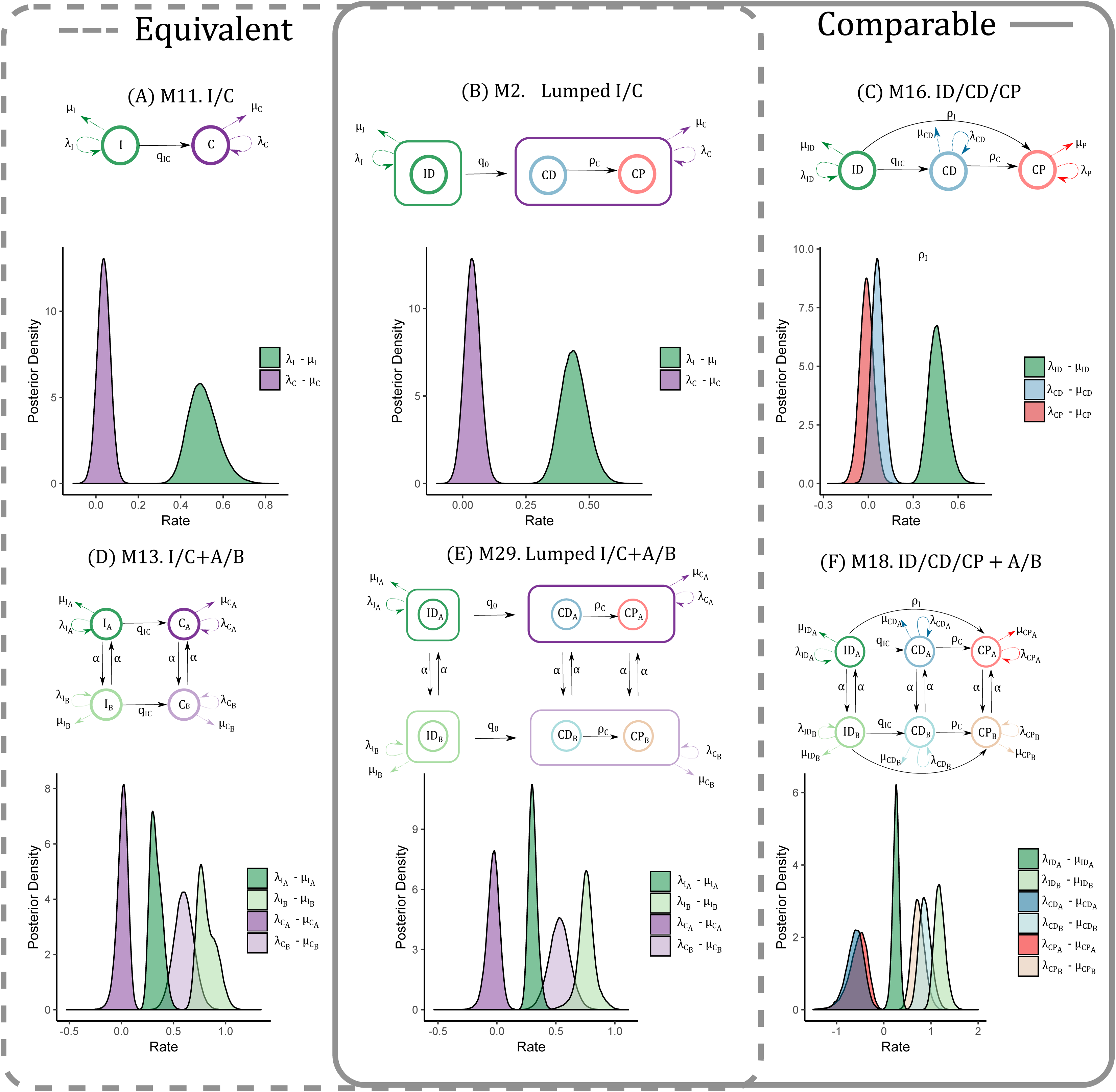
Lumped breeding system models to assess the effect of adding ploidy to create a three-state model. Moderate to insignificant evidence exists that adding ploidy is necessary Table 3

**Figure S5:**
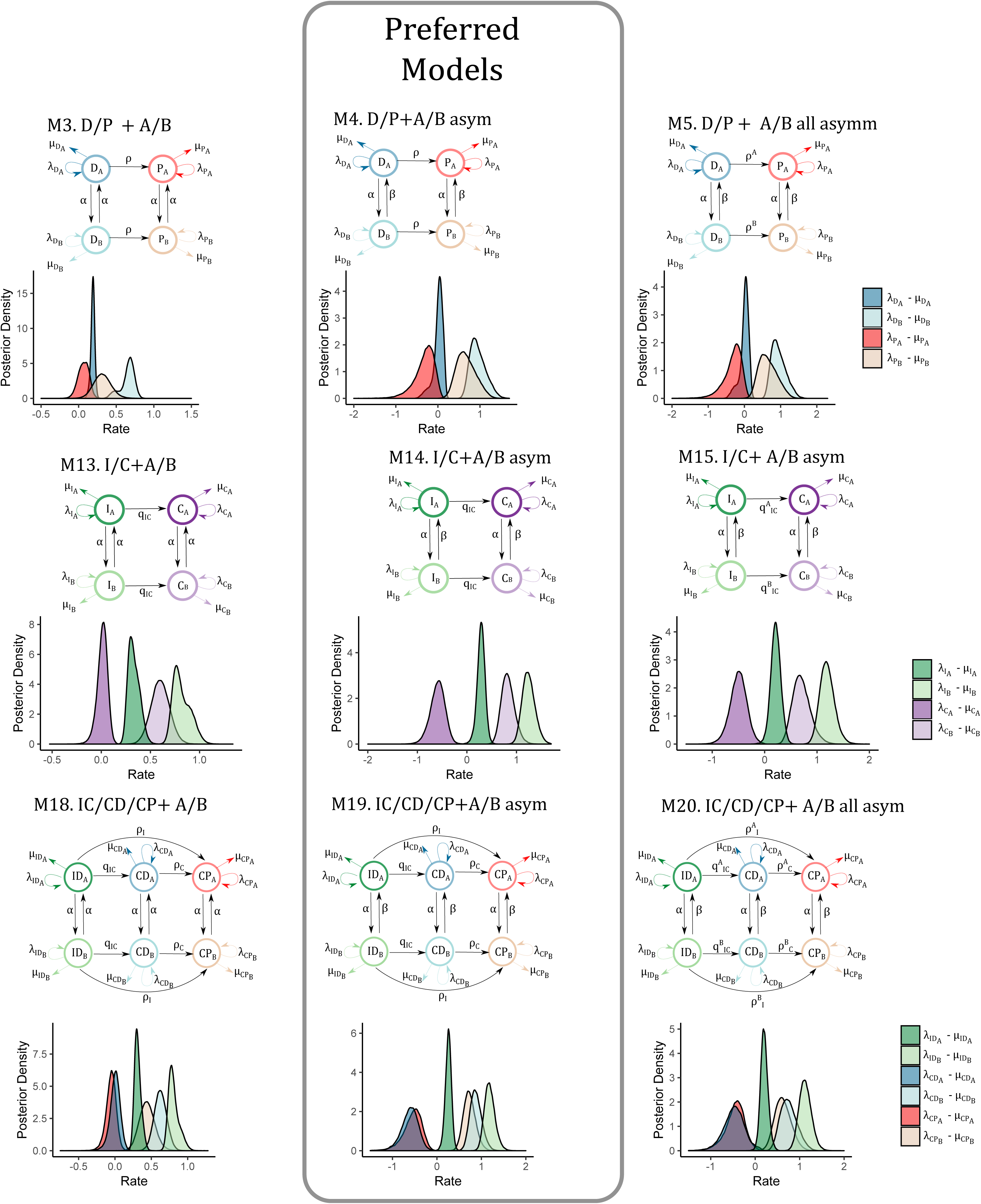
Effect of asymmetric rates in hidden models. Models with asymmetric rates are preferred over models with equal rates Table S6

**Figure S6:**
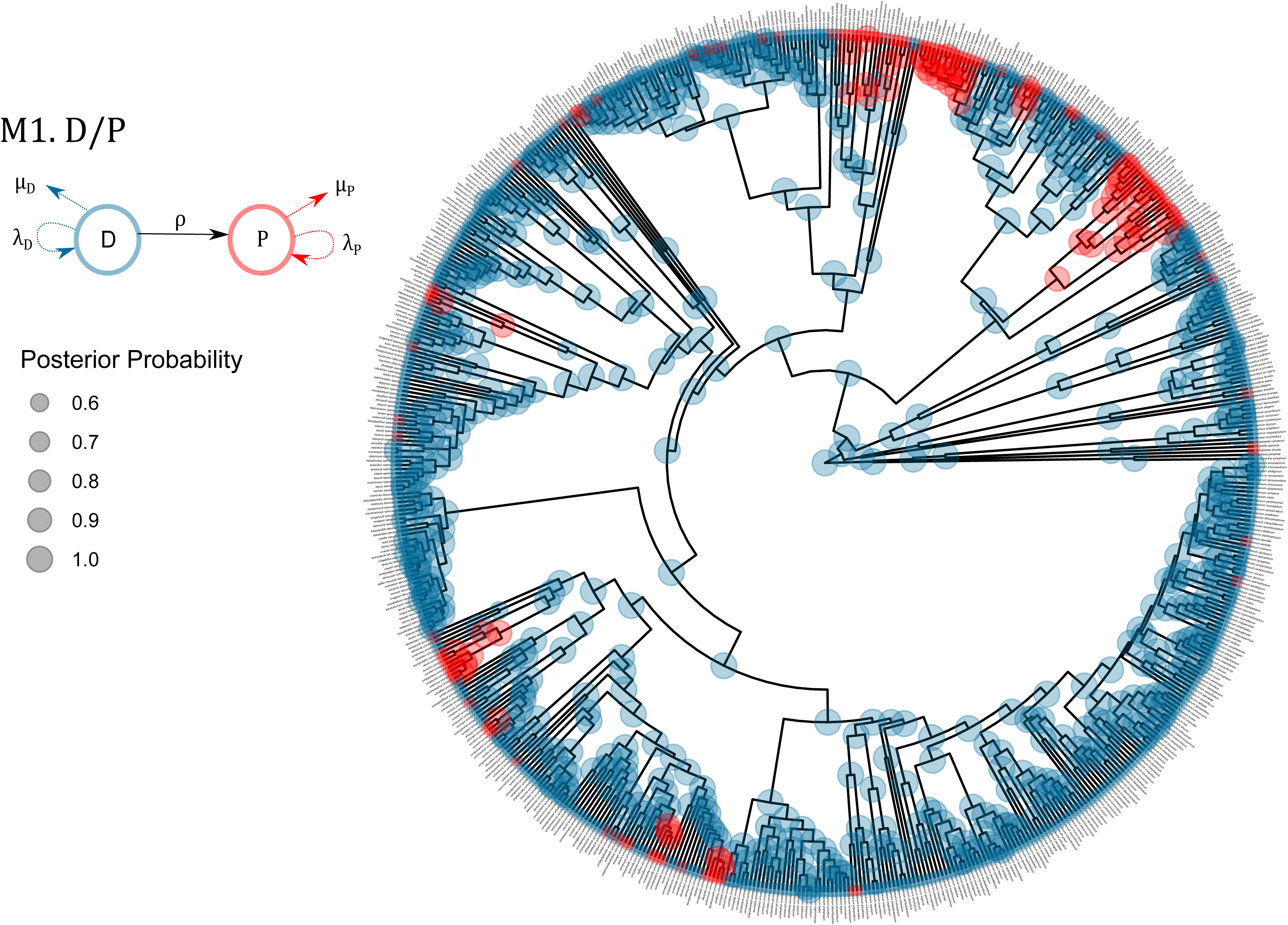
Ancestral state estimation using the maximum a posteriori for each node of theM1. D/P ploidy model

**Figure S7:**
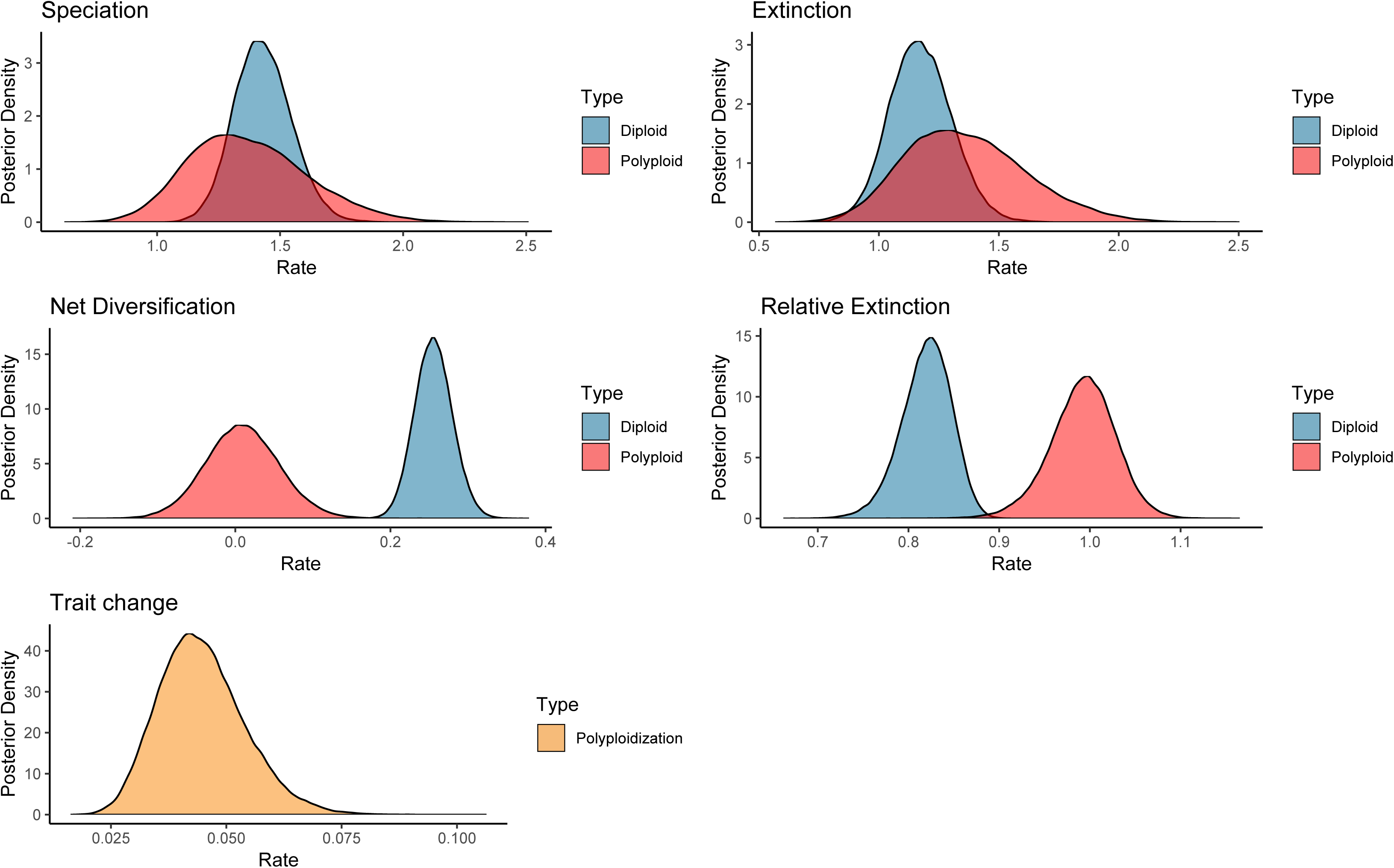
Posterior distribution for each of the parameters in the M1. D/P model

**Figure S8:**
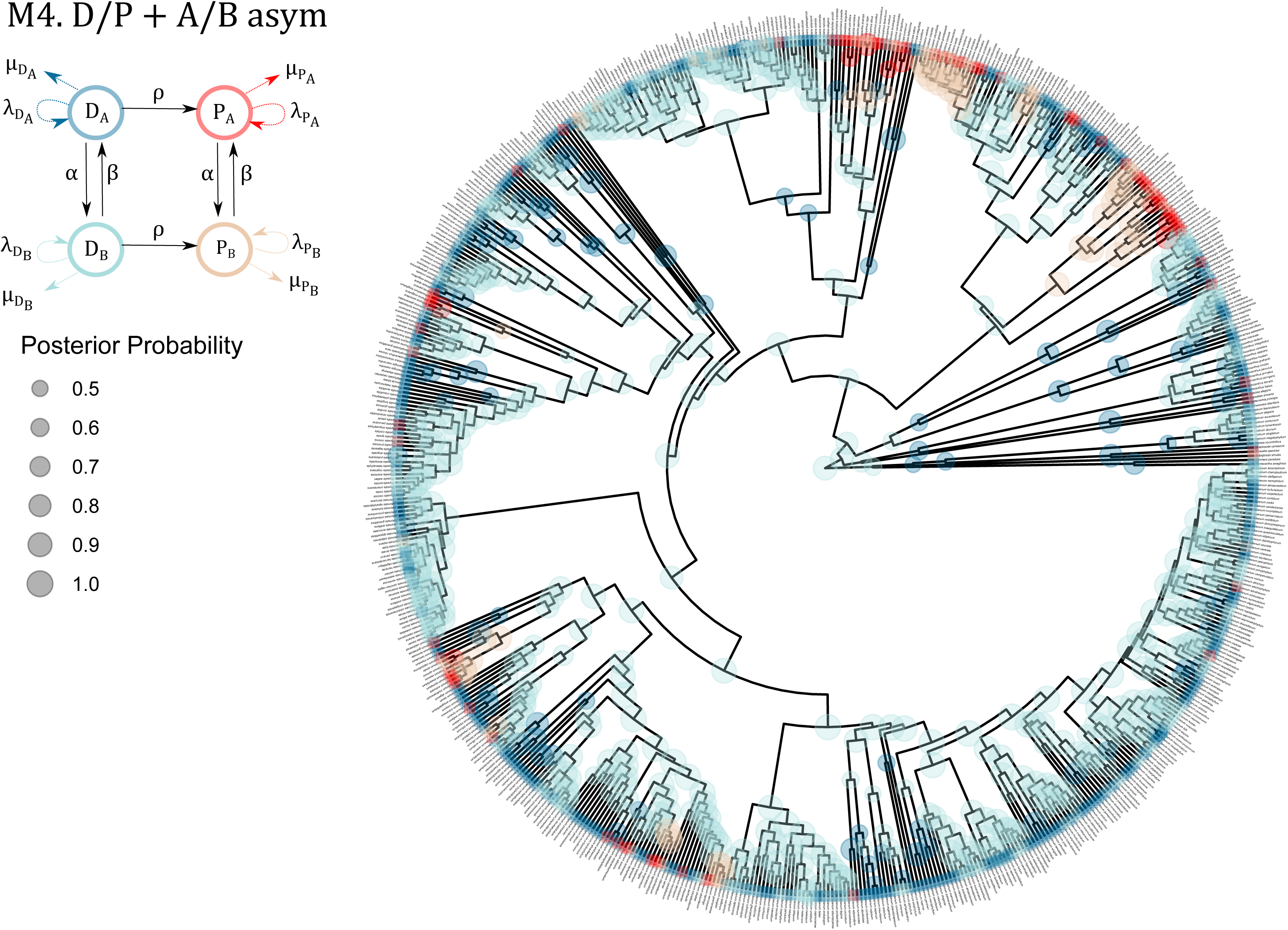
Ancestral state estimation using the maximum a posteriori for each node of the M4. D/P+A/B asym model

**Figure S9:**
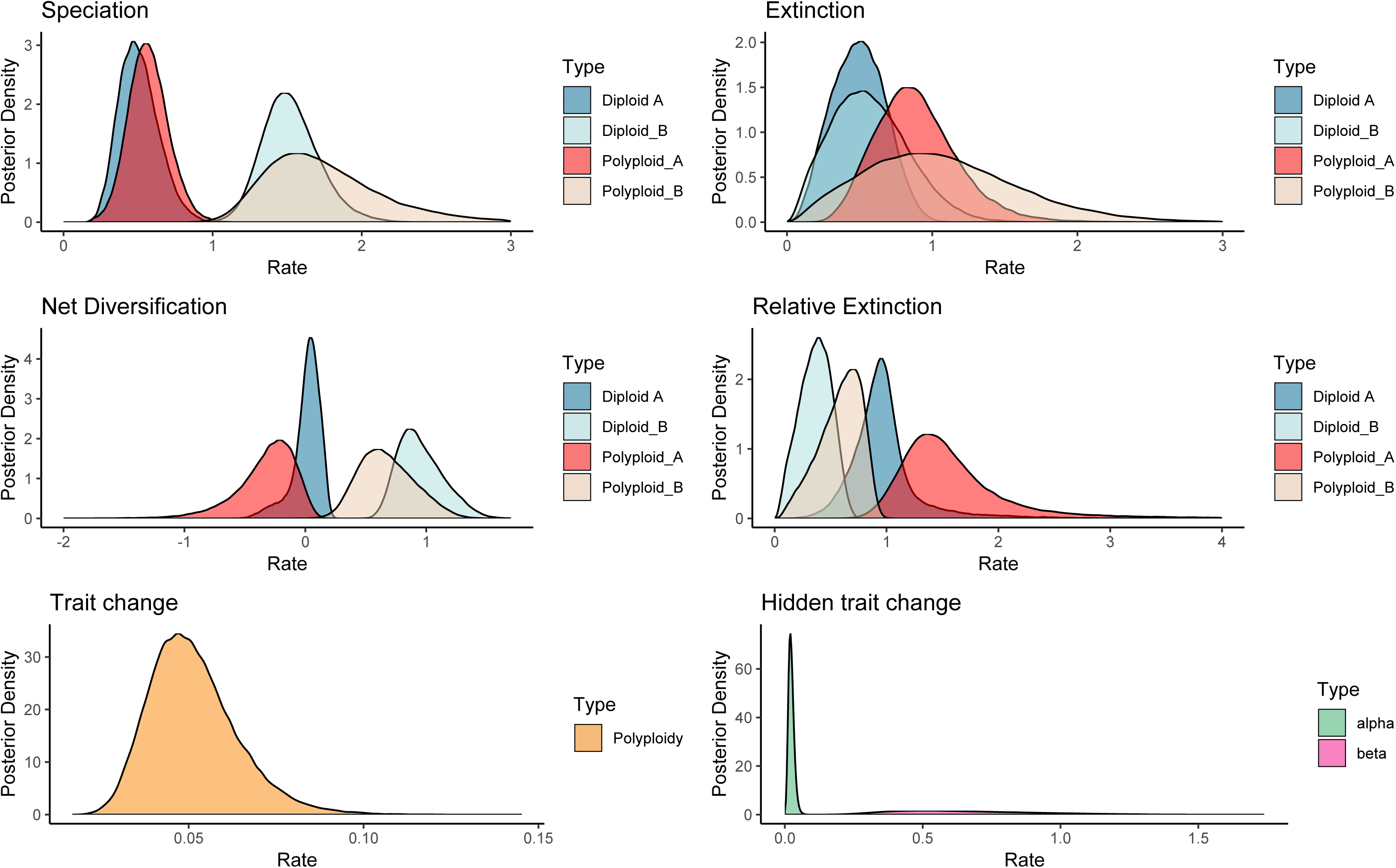
Posterior distribution for each of the parameters in the M4. D/P+A/B asym model

**Figure S10:**
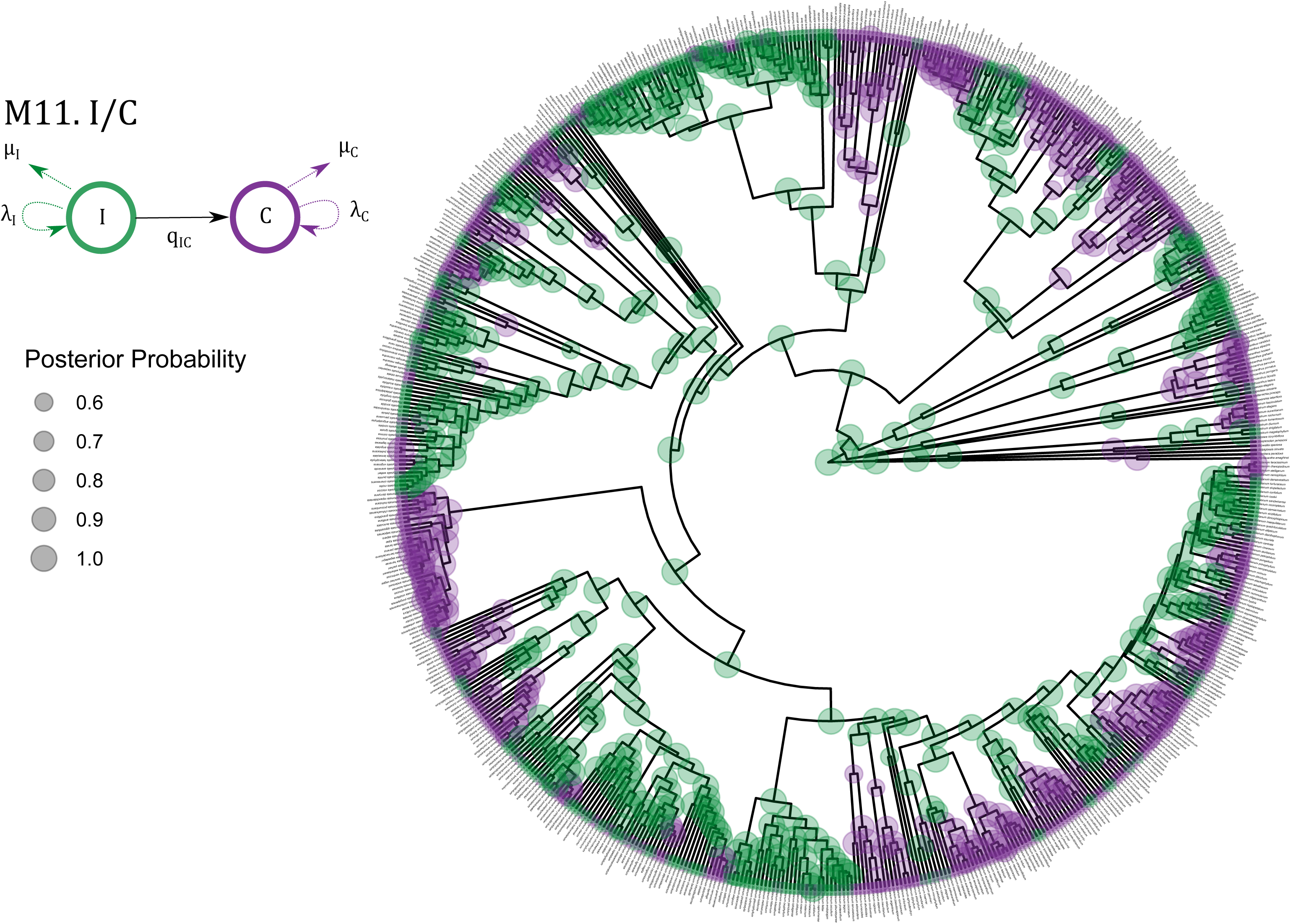
Ancestral state estimation using the maximum a posteriori for each node of the M11.I/C model

**Figure S11:**
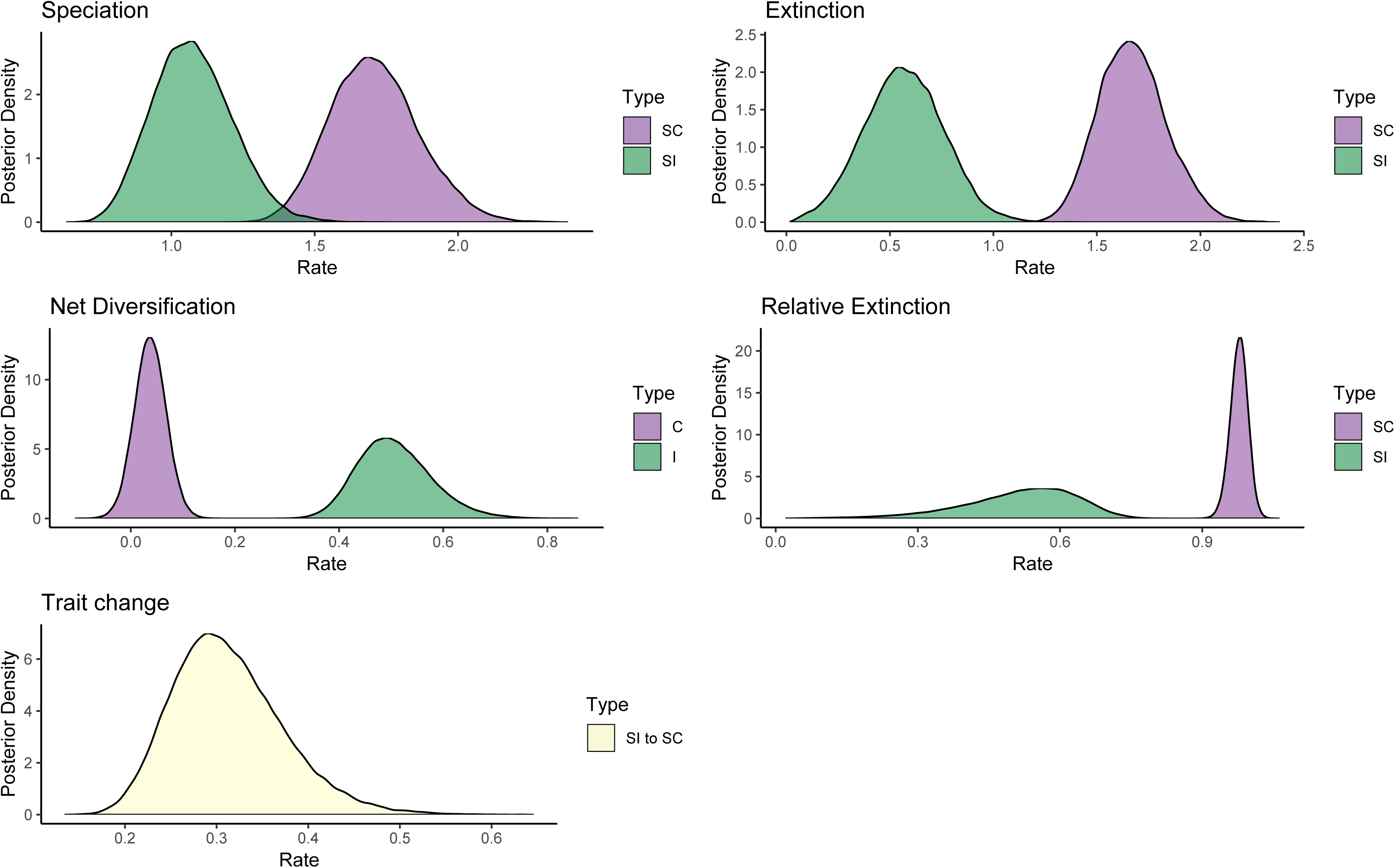
Posterior distribution for each of the parameters in the M11. I/C model

**Figure S12:**
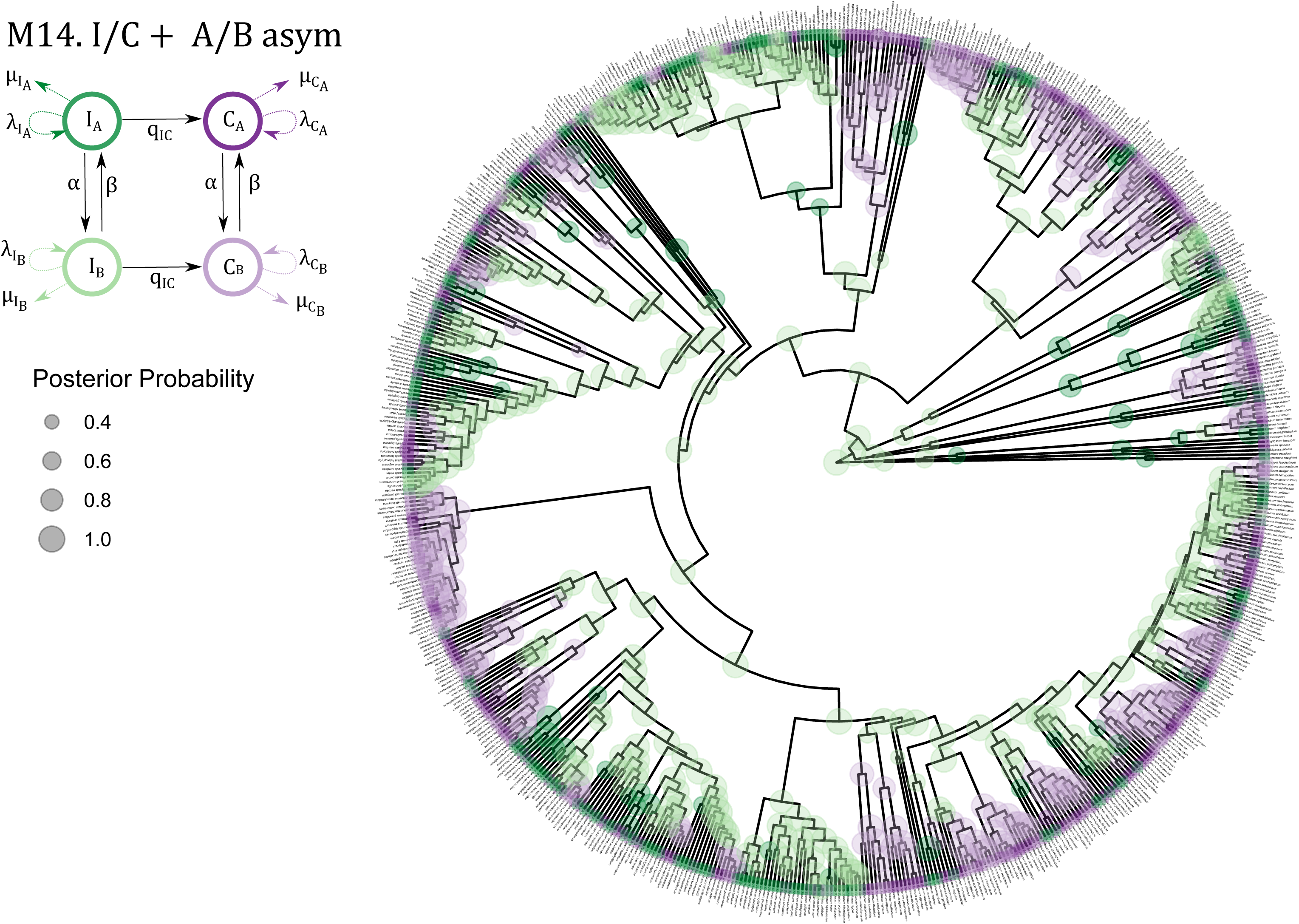
Ancestral state estimation using the maximum a posteriori for each node of the M14. I/C+A/B asym model

**Figure S13:**
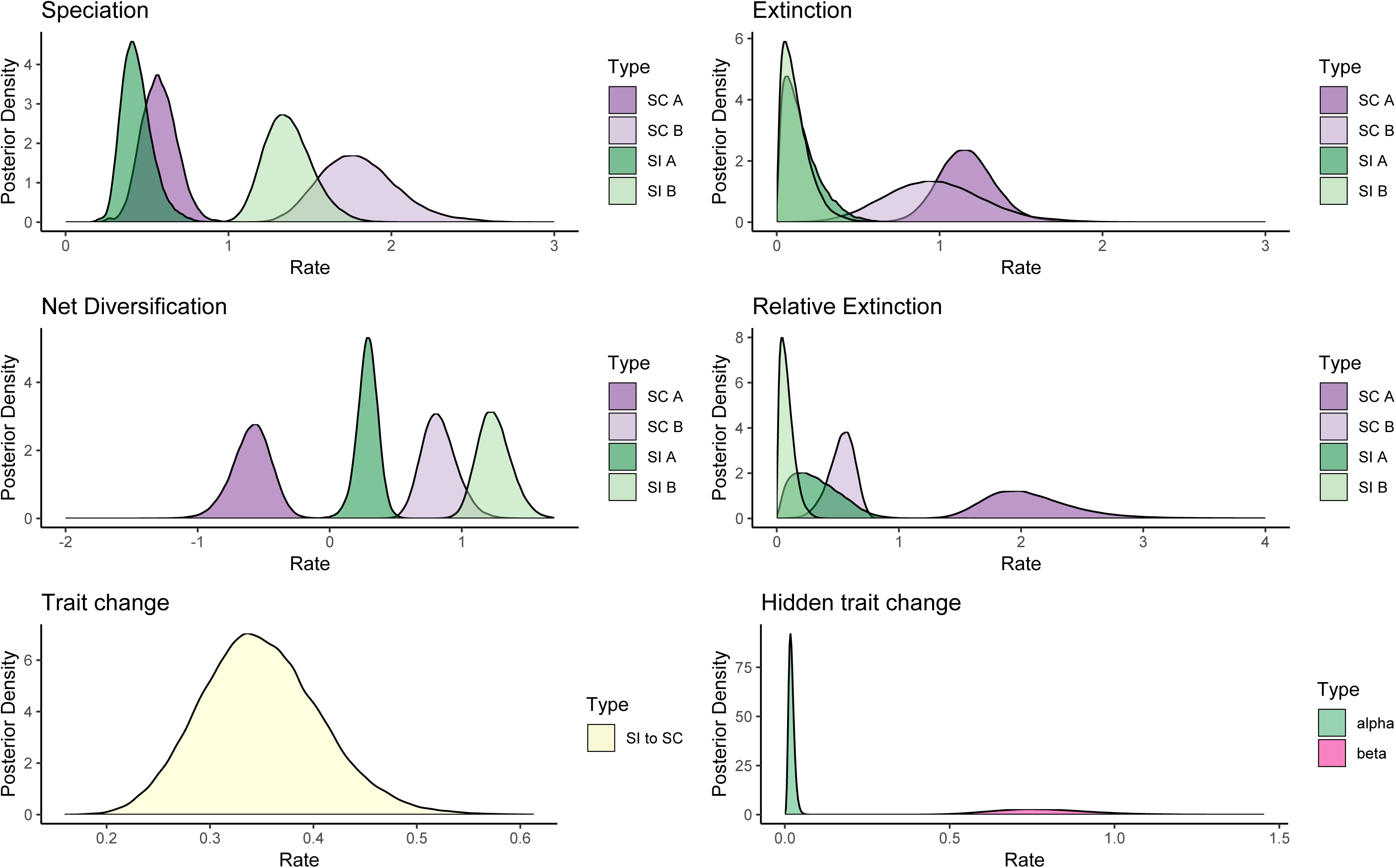
Posterior distribution for each of the parameters in the M14. I/C+A/B asym model

**Figure S14:**
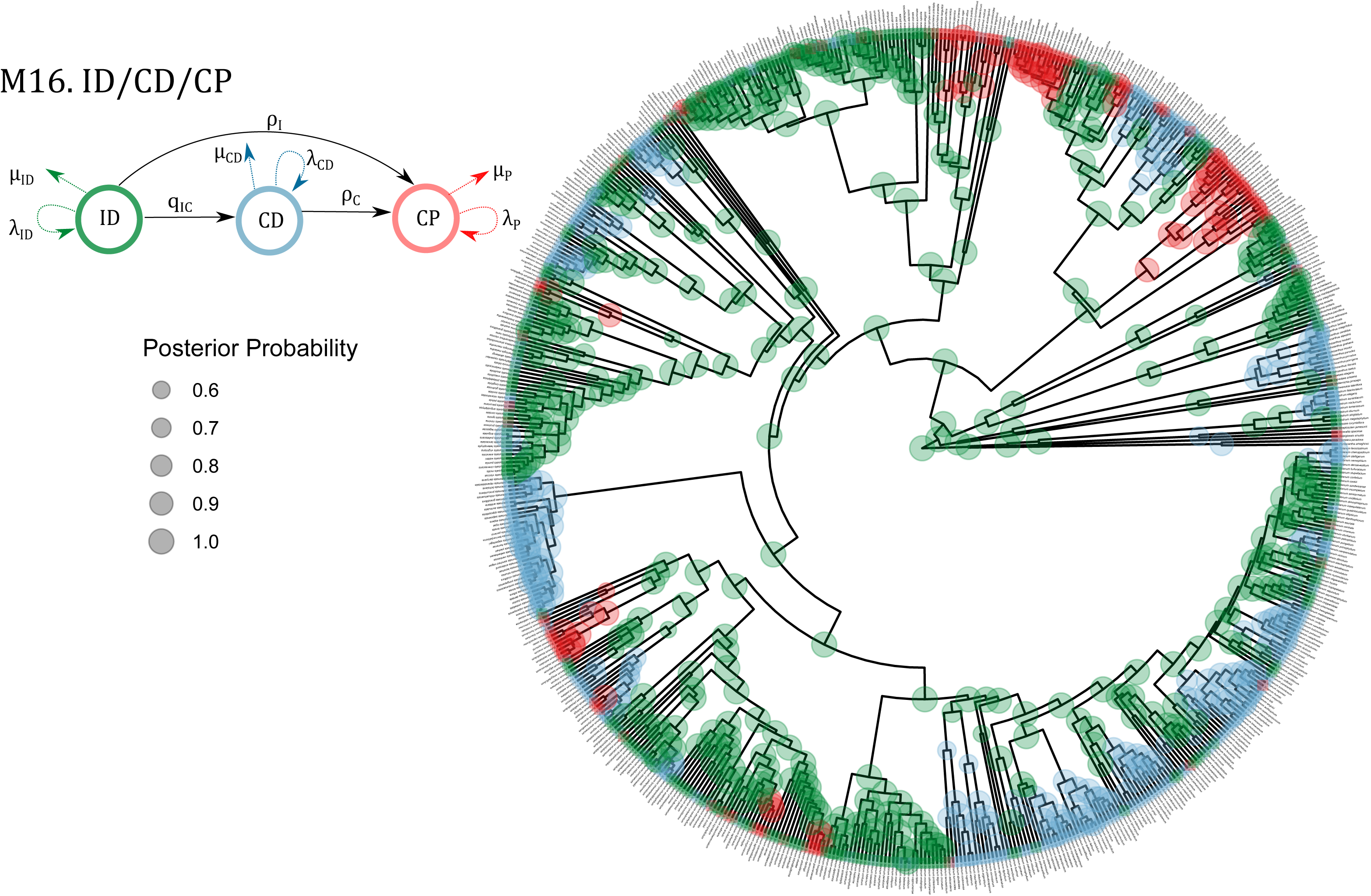
Ancestral state estimation using the maximum a posteriori for each node of the M16. ID/CD/CP model

**Figure S15:**
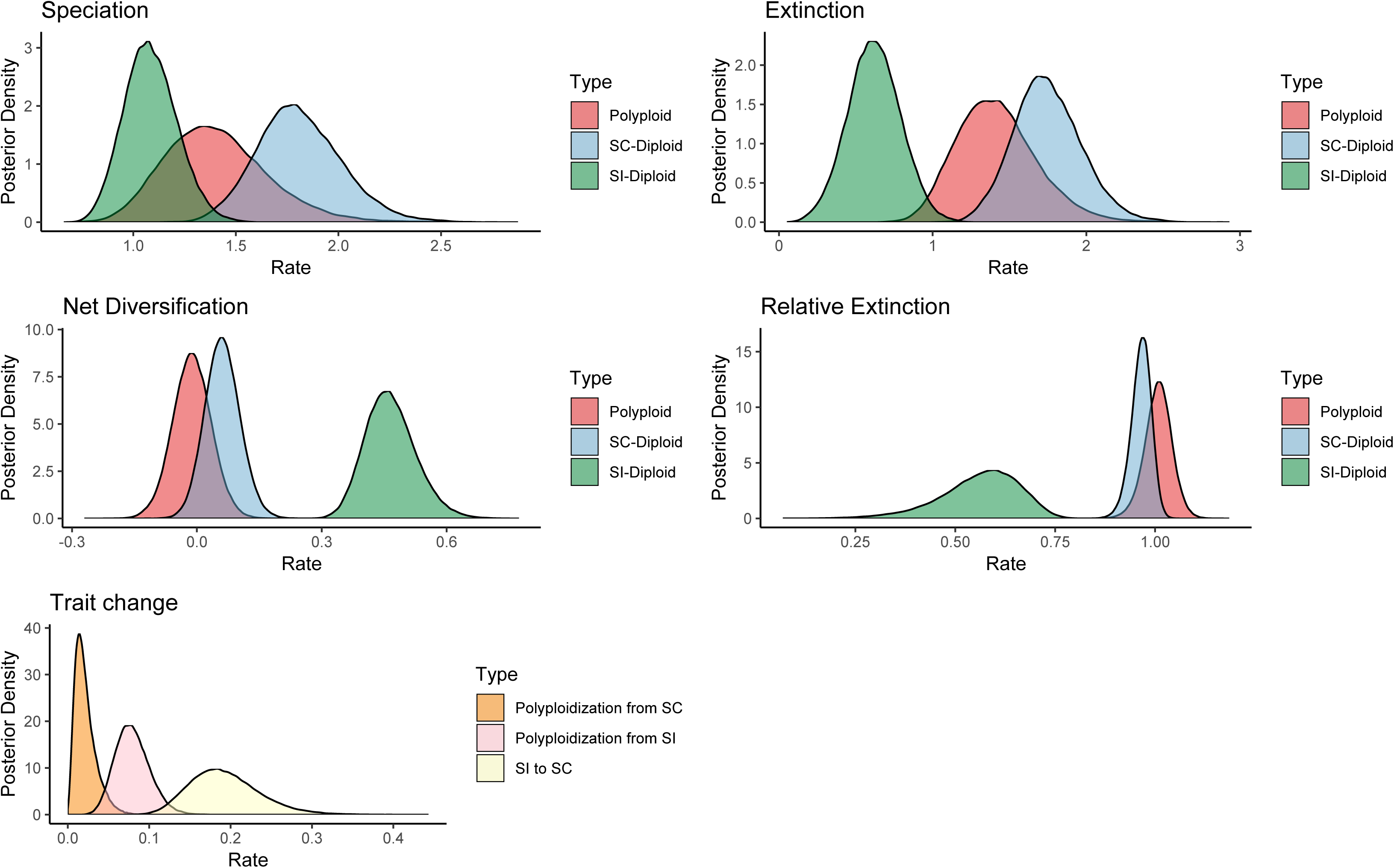
Posterior distribution for each of the parameters in the M16. ID/CD/CP model

**Figure S16:**
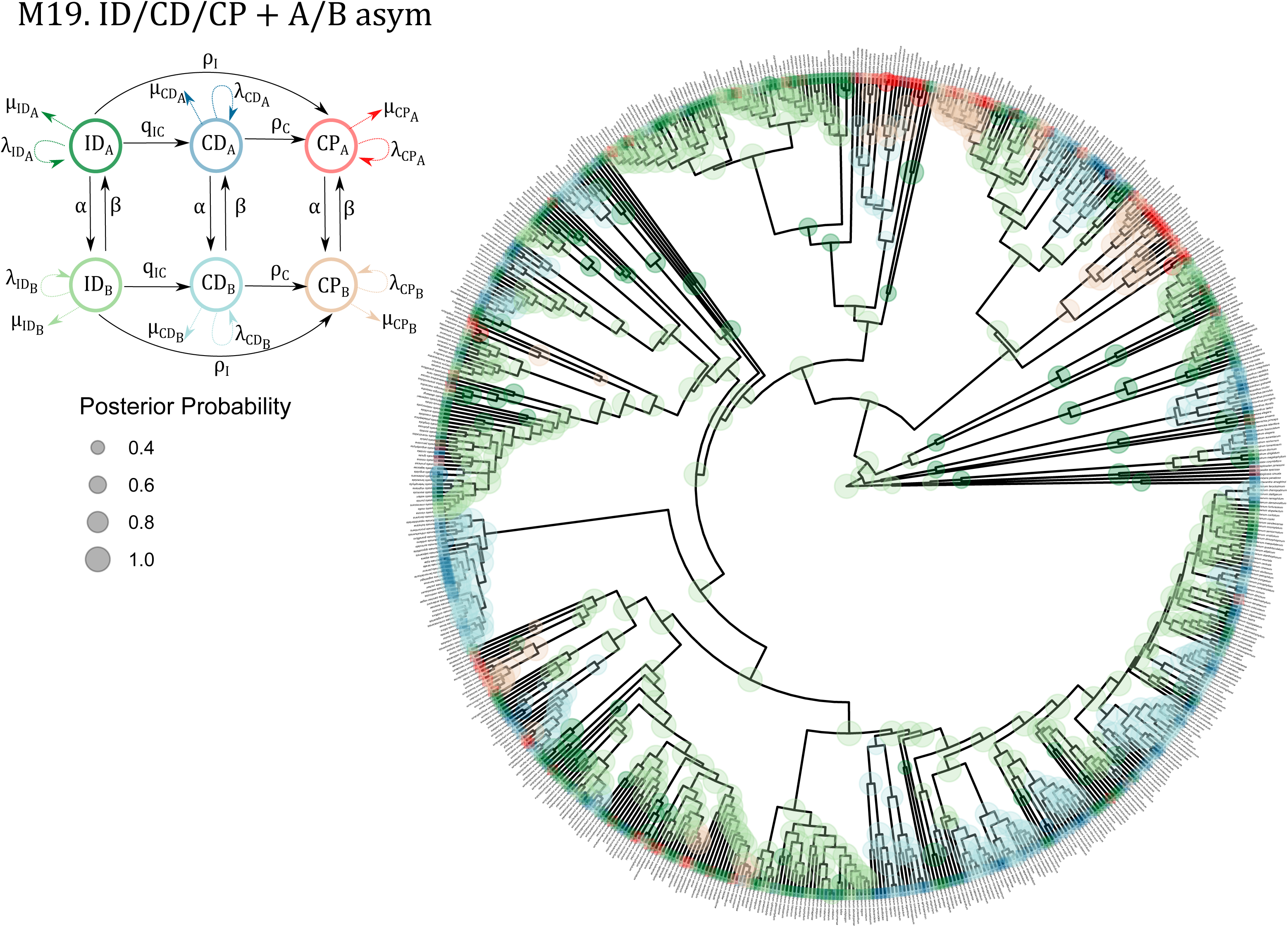
Ancestral state estimation using the maximum a posteriori for each node of the M19. ID/CD/CP+A/B asym model

**Figure S17:**
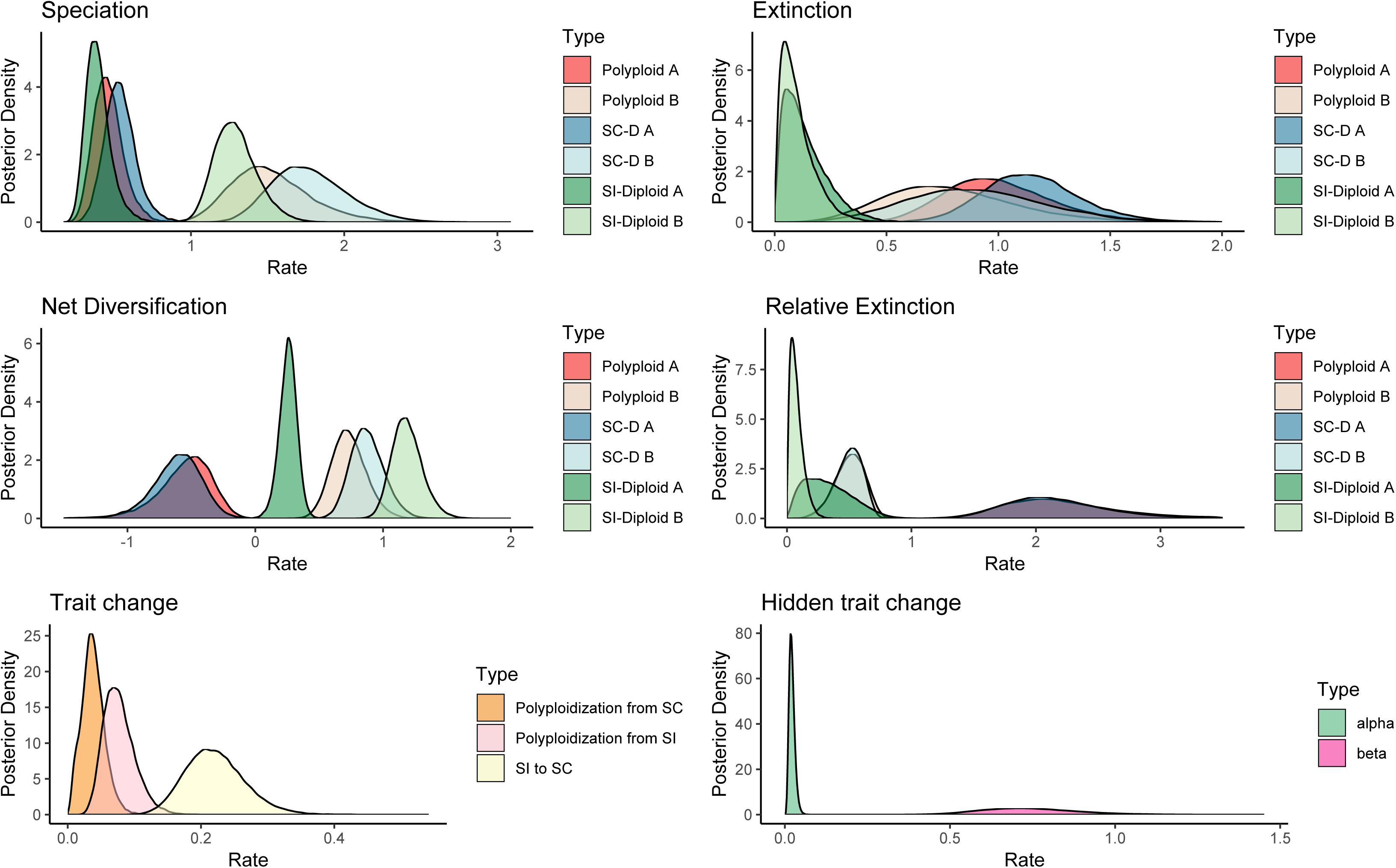
Posterior distribution for each of the parameters in the M19. ID/CD/CP+A/B asym model

**Table S1:**
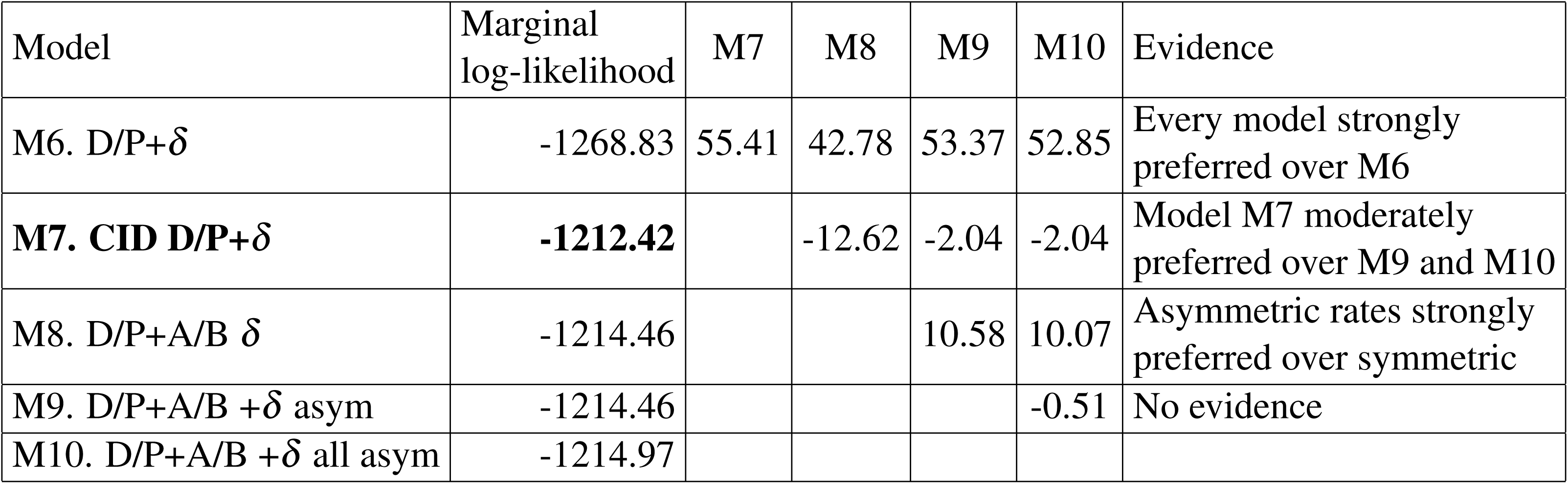
Bayes factors of ploidy-only models with diplodization. Results indicate that a character independent model (M6) is strongly preferred over BiSSE (M1) model. M6 is moderately preferred over any of the HiSSE models with asymmetric hidden rates (M9, M10).

| Model | Marginal<br>log-likelihood | M7 | M8 | M9 | M10 | Evidence |
| --- | --- | --- | --- | --- | --- | --- |
| M6. D/P+$\delta$ | -1268.83 | 55.41 | 42.78 | 53.37 | 52.85 | Every model strongly preferred over M6 |
| **M7. CID D/P+$\delta$** | **-1212.42** |  | -12.62 | -2.04 | -2.04 | Model M7 moderately preferred over M9 and M10 |
| M8. D/P+A/B $\delta$ | -1214.46 | | | 10.58 | 10.07 | Asymmetric rates strongly preferred over symmetric |
| M9. D/P+A/B +$\delta$ asym | -1214.46 | | | | -0.51 | No evidence |
| M10. D/P+A/B +$\delta$ all asym | -1214.97 | | | | | |

**Table S2:** Bayes factors of ploidy and breeding system without diploidization. Results indicate that the MuHiSSE model with asymmetric hidden rates (17) is strongly preferred over M16 and M18 and moderately preferred over the MuHiSSe with all rates asymetric (M20). *Marginal log-likelihood for M17 could not be calculated within allotted computer time.

| Model | Marginal log-likelihood | M18 | M19 | M20 | Evidence |
| --- | --- | --- | --- | --- | --- |
| M16. ID/CP/CD | -1459.11 | 45.11 | 65.91 | 63.98 | Every model strongly preferred over M16 |
| M17. CID ID/CP/CD | * |  |  |  |  |
| M18. ID/CP/CD+A/B | -1414 |  | 20.79 | 18.87 | Asymmetric rates strongly preferred over symmetric |
| **M19. ID/CP/CD +A/B asym** | **-1393.20** |  |  | -1.92 | Asymmetric hidden rates moderately preferred over all asymmetric |
| M20. ID/CP/CD +A/B all asym | -1393.12 |  |  |  |  |

**Table S3:** Bayes factors of ploidy and breeding system with diploidization. Results indicate that the MuHiSSE model with asymmetric hidden rates (M24) is strongly preferred over M21-M23 and moderately preferred over the MuHiSSe with all rates asymmetric (M25).

| Model | Marginal log-likelihood | M22 | M23 | M24 | M25 | Evidence |
| --- | --- | --- | --- | --- | --- | --- |
| M21. ID/CP/CD+$\delta$ | -1454.68 | 55.48 | 46.031 | 67.94 | 65.15 | Every model strongly preferred over M21 |
| M22. CID ID/CP/CD+$\delta$ | -1399.201 | | -9.452 | 12.45 | 9.675 | Model M24 strongly preferred over M22 |
| M23. ID/CP/CD+A/B $\delta$ | -1408.65 | | | 21.91 | 19.12 | Asymmetric rates strongly preferred over symmetric |
| **M24. ID/CP/CD +$\delta$ asym** | **-1386.74** |  |  |  | -2.78 | Asymmetric hidden rates preferred over all asymmetric |
| M25. ID/CP/CD +$\delta$ all asym | -1389.52 | | | | | |

**Table S4:**
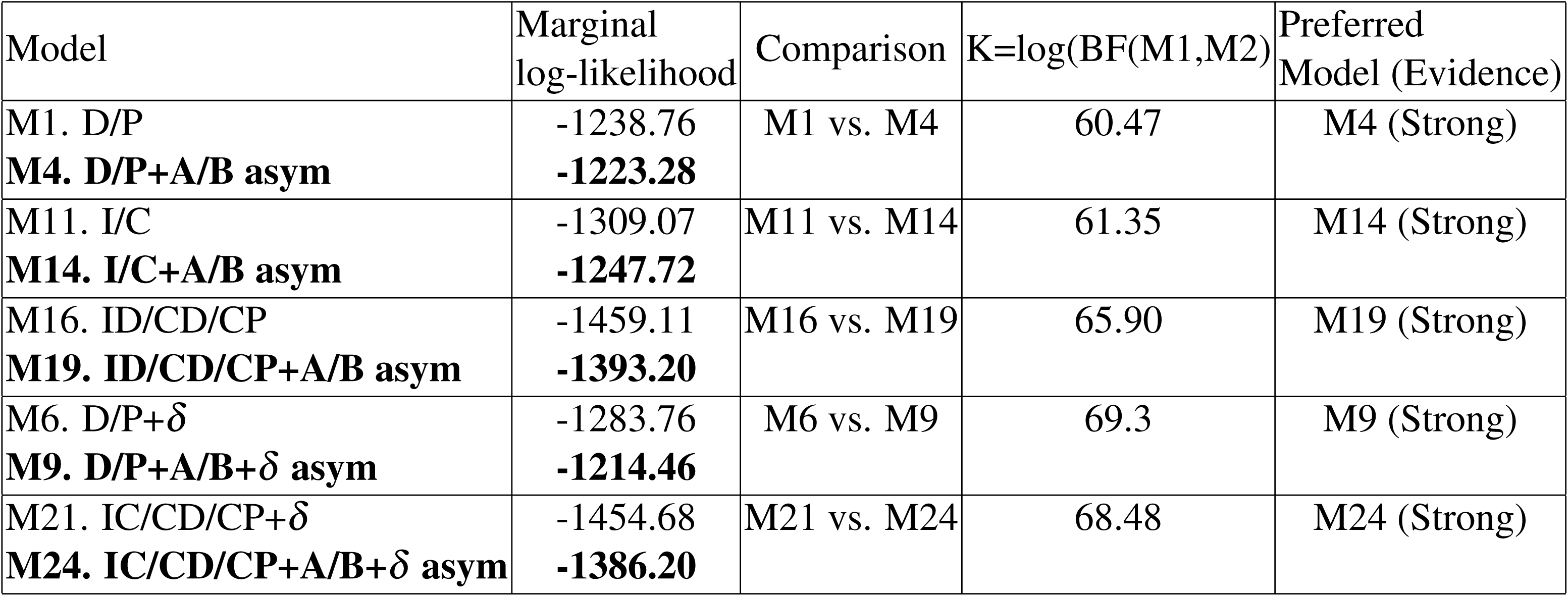
Test for addition of hidden states in models via Bayes factors. Models with hidden states are strongly preferred over simpler models

| Model | Marginal<br>log-likelihood | Comparison | $K=\log(\text{BF}(\text{M1}, \text{M2}))$ | Preferred<br>Model (Evidence) |
| --- | --- | --- | --- | --- |
| M1. D/P<br>**M4. D/P+A/B asym** | -1238.76<br>**-1223.28** | M1 vs. M4 | 60.47 | M4 (Strong) |
| M11. I/C<br>**M14. I/C+A/B asym** | -1309.07<br>**-1247.72** | M11 vs. M14 | 61.35 | M14 (Strong) |
| M16. ID/CD/CP<br>**M19. ID/CD/CP+A/B asym** | -1459.11<br>**-1393.20** | M16 vs. M19 | 65.90 | M19 (Strong) |
| M6. D/P+$\delta$<br>**M9. D/P+A/B+$\delta$ asym** | -1283.76<br>**-1214.46** | M6 vs. M9 | 69.3 | M9 (Strong) |
| M21. IC/CD/CP+$\delta$<br>**M24. IC/CD/CP+A/B+$\delta$ asym** | -1454.68<br>**-1386.20** | M21 vs. M24 | 68.48 | M24 (Strong) |

**Table S5:** Test for inclusion of a diploidization rate via Bayes factors. Models with diploidization are moderately preferred over models that do not include a diploidization rate

| Model | Marginal<br>log-likelihood | Comparison | $K=\log(\text{BF}(\text{M1}, \text{M2}))$ | Preferred<br>Model (Evidence) |
| --- | --- | --- | --- | --- |
| M1. D/P<br>**M6. D/P+$\delta$** | -1238.76<br>**-1267.84** | M1 vs. M6 | 65.92 | M6 (Strong) |
| M4. D/P+A/B asym<br>**M9. D/P+A/B+$\delta$ asym** | -1223.28<br>**-1214.46** | M4 vs. M9 | 8.81 | M9 (Moderate) |
| M16. ID/CD/CP<br>**M21. ID/CD/CP+$\delta$** | -1459.11<br>**-1454.68** | M16 vs. M21 | 4.41 | M21 (Moderate) |
| M19. IC/CD/CP +A/B asym<br>**M24. IC/CD/CP+A/B+$\delta$ asym** | -1393.20<br>**-1386.20** | M19 vs. M24 | 6.46 | M24 (Moderate) |

**Table S6:** Test for symmetry of the hidden and trait rates via Bayes factors. For all models asymmetric hidden transition rates are preferred over models with equal hidden rates. Adding more complexity by assuming asymmetry in the trait rate within hidden states does not improve models.

| Model | Marginal<br>log-likelihood | Comparison | K=log(BF(M1,M2)) | Preferred<br>Model (Evidence) |
| --- | --- | --- | --- | --- |
| M3. D/P+A/B | -1234.52 | M3 vs. M4 | 11.239 | M4 (Strong) |
| **M4. D/P+ A/B asym** | -1223.28 | M4 vs. M5 | -1.658 | M4 (Moderate) |
| M5. D/P+A/B all asym | -1224.93 |  |  |  |
| M13. I/C+ A/B | -1270.47 | M13 vs. M14 | 22.75 | M14 (Strong) |
| **M14. I/C+ A/B asym** | **-1247.72** | M14 vs. M15 | 0.05 | No evidence |
| M15. I/C+ A/B all asym | -1247.67 |  |  |  |
| M18. IC/CD/CP+A/B | -1414.00 | M18 vs. M19 | 20.79 | M19 (Strong) |
| **M19. IC/CD/CP+ A/B asym** | **-1393.21** | M19 vs. M20 | -1.919 | M19 (Moderate) |
| M20. IC/CD/CP+ A/B all asym | -1395.129 |  |  |  |
| M8. D/P +A/B+$\delta$ | -1225.05 | M8 vs. M9 | 10.58 | M9 (Strong) |
| **M9. D/P+ A/B+ $\delta$ asym** | **-1214.46** | M9 vs. M10 | -0.52 | M10 (Moderate) |
| M10. D/P+A/B+$\delta$ all asym | -1214.98 | | | |
| M23. IC/CD/DP+A/B+$\delta$ | -1408.65 | M23 vs M24 | 21.91 | M24(Strong) |
| **M24. IC/CD/DP+A/B+$\delta$ asym** | **-1386.74** | M24 vs M25 | -2.78 | M24 (Moderate) |
| M25. IC/CD/DP+A/B+ $\delta$ all asym | -1389.52 | | | |

